# Design and off-target prediction for antisense oligomers targeting bacterial mRNAs with the MASON webserver

**DOI:** 10.1101/2022.05.24.492283

**Authors:** Jakob Jung, Linda Popella, Phuong Thao Do, Patrick Pfau, Jörg Vogel, Lars Barquist

**Affiliations:** Institute for Molecular Infection Biology, University of Würzburg, Würzburg, Germany; Helmholtz Institute for RNA-based Infection Research (HIRI), Helmholtz Centre for Infection Research (HZI), Würzburg, Germany; Faculty of Medicine, University of Würzburg, Würzburg, Germany

**Keywords:** peptide nucleic acid, antisense oligomers, antisense antibiotics, RNA Sequencing, webserver, off-targets

## Abstract

Antisense oligomers (ASOs) such as peptide nucleic acids (PNAs), designed to inhibit the translation of essential bacterial genes, have emerged as attractive sequence- and species-specific programmable RNA antibiotics. Yet, potential drawbacks include unwanted side effects caused by their binding to transcripts other than the intended target. To facilitate the design of PNAs with minimal off-target effects, we developed MASON (**M**ake **A**nti**S**ense **O**ligomers **N**ow), a webserver for the design of PNAs that target bacterial mRNAs. MASON generates PNA sequences complementary to the translational start site of a bacterial gene of interest and reports critical sequence attributes and potential off-target sites. We based MASON’s off-target predictions on experiments in which we treated *Salmonella enterica* serovar Typhimurium with a series of 10mer PNAs derived from a PNA targeting the essential gene *acpP* but carrying two serial mismatches. Growth inhibition and RNA-sequencing (RNA-seq) data revealed that PNAs with terminal mismatches are still able to target *acpP*, suggesting wider off-target effects than anticipated. Comparison of these results to an RNA-seq dataset from uropathogenic *Escherichia coli* (UPEC) treated with eleven different PNAs confirmed our findings are not unique to *Salmonella*. We believe that MASON’s off-target assessment will improve the design of specific PNAs and other ASOs.

## INTRODUCTION

Antimicrobial resistant (AMR) bacteria have become a major threat to human health, underscoring the urgency for the development of new types of antibiotics (Murray et al. 2022). Most clinically available antibiotics disrupt conserved cellular processes like transcription, translation, DNA replication, or cell wall maintenance (reviewed in (Baquero and Levin 2021)). Therefore, antibiotic treatment often affects a wide spectrum of microbes and can disturb the natural balance of the human microbiome, potentially leading to dysbiosis (Vangay et al. 2015). To prevent this effect, ultra-narrow spectrum antibiotics targeting specific pathogenic species or strains would be an important advance in antimicrobial research (Vogel 2020).

A promising approach is the use of antisense oligomers (ASOs) that inhibit the translation of essential genes by binding to the translation initiation region (TIR), making them highly specific. Popular ASOs include peptide nucleic acids (PNAs), which are synthetic nucleic acid analogs commonly designed to bind the ribosome binding site (RBS) or start codon of a gene of interest (Dryselius et al. 2003; Nielsen 2006; Egholm et al. 1993; Nielsen et al. 1991). Their neutral pseudopeptide backbone protects PNAs from degradation by nucleases and proteases while conferring higher binding affinities compared to DNA or RNA. To facilitate cellular uptake by bacteria, PNAs are coupled to short (<30 aa), often positively charged cell penetrating peptides (CPPs) (Eriksson et al. 2002). PNAs have been successfully applied to inhibit growth of a wide variety of microorganisms both in culture and in animal models (Hegarty and Stewart 2018; Sully and Geller 2016; Barkowsky et al. 2019; Pifer and Greenberg 2020; Tan et al. 2005; Lee et al. 2019; Oh et al. 2014). The ability of PNAs to selectively silence any gene of interest can also have other applications, such as to potentiate conventional antibiotics by silencing nonessential genes involved in drug resistance or tolerance (Aunins et al. 2020; Courtney and Chatterjee 2015).

Despite nearly two decades of research, design rules for PNA sequences are based on only a handful of studies. Early work in *E. coli* investigating PNAs of varying lengths suggested that a length of 9 to 12 bases is optimal to reliably inhibit target gene translation (Good et al. 2001). More recently, PNA length was shown to be a major constraint for cell-entry, and 10 bases was proposed as an ideal length (Goltermann et al. 2019). Regarding sequence composition, PNAs rich in purines show reduced solubility (Nielsen et al. 1996; Good 2002), while a low GC content can result in low binding affinities between PNAs and its target (Giesen et al. 1998). Further, self-complementarity of PNA sequences needs to be avoided to prevent self-interaction (Good 2002).

Another important consideration for PNA design is target site uniqueness. Because of the short length of PNAs, full or partial complementarity to genomic regions other than target region is frequent. In addition, the TIR, spanning the Shine-Dalgarno sequence (SD) and/or start codon, is often not completely unique. We refer to these unintended complementary regions as off-target matches. Naturally, off-target matches with one or two base pair (bp) mismatches are very common for 10-mer PNAs in bacteria. It is thus important to determine whether mismatches interfere with PNA targeting. Cell-free hybridization experiments have established that single mismatches in PNA-DNA and PNA-RNA duplexes substantially decrease binding affinity (Ratilainen et al. 2000; Jensen et al. 1997). In addition, PNAs with single and double mismatches to bacterial target mRNAs have strongly reduced effects on target gene expression and growth inhibition compared to fully complementary PNAs (Hatamoto et al. 2009; Good et al. 2001; Good and Nielsen 1998). A more recent study, in which single mismatches were introduced in PNAs of different lengths targeting the *acpP* gene in *E. coli* showed a negative correlation between PNA melting temperatures and minimum inhibitory concentrations (Goltermann et al. 2019). However, these results were based on a small number of tested PNAs, making it difficult to draw general conclusions. Nevertheless, PNAs are thought to tolerate very few mismatches, making them highly specific nucleic acid-binding compounds.

In bacterial PNA design, little attention has therefore been given to off-targets that harbor one or two bp mismatches. Yet, broad consideration of all possible off-target effects is essential for several reasons. First, PNAs that bind to TIRs of other genes can cause unwanted cellular responses. This may be particularly problematic when targeting non-essential genes with the goal of selectively silencing gene expression rather than killing the bacteria. Second, binding to a large number of off-target sites may reduce PNA availability by sequestering free PNAs, a phenomenon known from bacterial small RNA sponges (Ziebuhr and Vogel 2015). While it has been generally recommended to avoid off-target matches during PNA design (Good 2002; Dryselius et al. 2003), the degree of complementarity required to mediate off-target effects as well as the overall transcriptomic response caused by off-targets have not been investigated in detail yet.

Although there are some guidelines for the design of effective PNA sequences (Dryselius et al. 2003), researchers usually rely on manual design to find the best sequence to target a selected gene. This is because the existing computational tools either lack a user-friendly interface, or are designed for ASOs targeting mammalian cells. *PNA Finder*, for example, can be used to design PNA sequences and screens for sequence-specific attributes and off-target effects (Eller et al. 2021). However, because it requires installation of dependencies in a Bash shell, it may be difficult to use for those with limited computational experience. Other methods, such as *PNA tool* (https://www.pnabio.com/support/PNA_Tool.htm), are easy to use, but only screen for sequence-specific attributes, without considering off-target effects in bacteria.

Here, to facilitate the design of bacterial ASOs we developed MASON (**M**ake **A**nti**S**ense **O**ligomers Now), a user-friendly webserver which helps researchers in designing PNAs for any annotated bacterial gene while predicting melting temperature (T_m_) and possible off-targets in the bacterium of interest. To improve the off-target prediction algorithm of MASON and to investigate the effect of mismatches on PNA efficiency, we designed a series of 10mer PNAs with serial double mismatches based on a PNA targeting the *acpP* mRNA of *Salmonella enterica* serovar Typhimurium strain SL1344 (henceforth *Salmonella*). We found that double mismatches at the termini of PNAs do not fully abrogate the growth inhibitory and transcript-depleting effects of a PNA. Accordingly, PNAs with partial complementarity in proximity to the translational start region of off-target mRNAs can substantially reduce transcript levels in both *Salmonella* and uropathogenic *Escherichia coli* (UPEC). We incorporated these findings into MASON to improve its off-target predictions. MASON is freely available as both a webserver and command line tool.

## RESULTS

### MASON - a webserver for efficient ASO design

To facilitate the design of bacterial ASOs, we developed a user-friendly web application, called MASON (https://www.helmholtz-hiri.de/en/datasets/mason, Supplemental Fig. S1). Users can specify the organism of interest, a target gene, the ASO length and the number of allowed mismatches for off-targets (Fig. 1A, Supplemental Fig. S2). A pre-defined set of four annotated bacterial strains, specifically *E. coli* str. K12 substr. MG1655, *Salmonella enterica* subsp. *enterica* serovar Typhimurium SL1344, *Clostridium difficile* 630 and *Fusobacterium nucleatum* ATCC 23726, can be selected from a dropdown menu (Supplemental Fig. S2). Furthermore, custom bacterial genomes can be added manually with a reference fasta file and an annotation file in gff format, which can be downloaded for a wide variety of bacteria from NCBI. Finally, there is an additional option to screen for off-targets in human genes and genes of the human microbiome.

**Figure 1.**
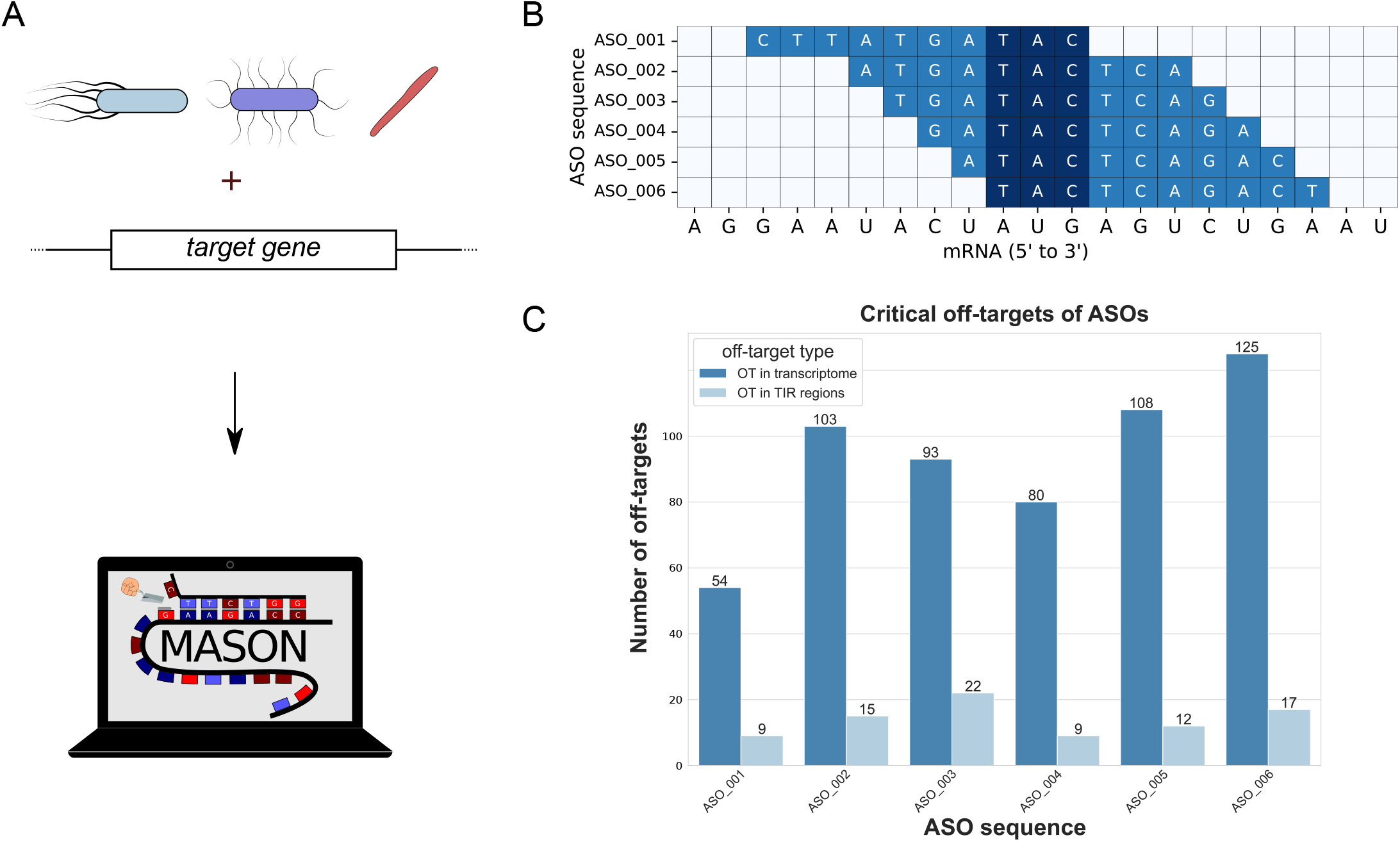
The MASON webserver designs ASO sequences and predicts off-targets. **(A)** Users specify an organism of interest, the target gene, ASO length and allowed number of mismatches for off-targets. MASON then generates ASO sequences and their attributes in between 5 and 30 seconds. **(B)** The generated ASO sequences are designed to bind the start codon (marked in dark blue) of the target mRNA. **(C)** MASON predicts critical off-target binding regions for each ASO. An off-target is termed “critical” if the respective ASO sequence binds more than 7 consecutive bases of the off-target mRNA. Off-targets are predicted both for the whole transcriptome (steel blue) and for translation initiation regions (TIR, light blue) of other genes. TIR off-targets are defined as having the start of the binding region between −20 to +5 bases relative to the start codon.

Once the user submits the input, MASON finishes its calculation in between ten seconds and two minutes, depending on the size of the target genome, choice of ASO length, and allowed number of mismatches for off-targets (Fig. 1A). For each selected target gene, MASON outputs all candidate ASO sequences that overlap the start codon (in a window ranging from −7 to +10 for 10mers, default) (Supplemental Fig. S3A). Because it is often desirable to target the RBS in the 5’ UTR of a transcript, the user can opt to specify the number of bases upstream of the start codon for which additional ASOs are designed.

Recent reports using different *E. coli* strains showed that PNA-mRNA T_m_ positively correlates with growth inhibition (Goltermann et al. 2019) and is associated with *in vitro* translation inhibition of the target gene (Popella et al. 2022). We therefore integrated T_m_ predictions for PNA-RNA duplexes as a feature in MASON (see methods, Supplemental Fig. S3B). To predict the T_m_, we applied a nearest-neighbor method for RNA-RNA duplexes (Xia et al. 1998), which is incorporated in the MELTING 5 platform (Dumousseau et al. 2012). Next, to test whether our predictions perform well, we compared them to experimental T_m_ measurements of PNA-RNA duplexes (Goltermann et al. 2019) and to a PNA-DNA prediction algorithm (Giesen et al. 1998) used by the *PNA tool* and the *PNA finder* toolbox (Eller et al. 2021) (Supplemental Fig. S4). Our predictions correlate well with the experimentally measured T_m_, but the absolute T_m_ was consistently underestimated by around 10 °C. This is likely due to increased stability of PNA-RNA duplexes compared to RNA-RNA interactions (Egholm et al. 1993). Interestingly, our predictions based on an RNA hybridization model were closer to experimental PNA-RNA stability measurements than those predicted by the PNA-DNA interaction prediction algorithm (Supplemental Fig. S4).

Due to the short length of ASOs, the frequency of off-target sites with zero or few mismatches can be high. We therefore established an off-target prediction algorithm in MASON. The algorithm screens for all possible off-target matches with a user-adjusted number of mismatches in the target organism’s annotated coding sequence (CDS) and 30 bases upstream (5’) of the respective CDS (see methods for details). We divide off-targets into two categories: (i) off-target sites in the whole transcriptome based on annotated CDSs; and (ii) off-target site in the TIR of non-target genes with start positions 20 bases upstream of the annotated start codon until 5 bases downstream (Fig. 1C, Supplemental Fig. S3C). The results page of MASON additionally provides a table that summarizes sequence properties, the number of self-complementary bases and off-target sites (Supplemental Fig. S5).

We also created a command line version of MASON, with added features that facilitate high-throughput design of ASOs. The command line version allows users to skip the design step and use previously designed sequences as direct input, thereby permitting parallel screening of a large number of ASO sequences in a short time. It also adds the possibility to screen multiple bacterial species for off-targets, which can be useful for applying ASOs to co-cultures or microbial communities. As we want to keep the web-interface fast and efficient, these functions are only available in the command line version of MASON, which runs on Linux-based PCs, servers or high-performance clusters.

### PNAs targeting acpP retain growth-inhibitory activity despite terminal double mismatches

The algorithm of MASON considers off-target matches only if at least seven consecutive matching bases exist. To investigate whether this definition of ‘critical’ off-targets is reasonable, we took a scanning approach to design a series of 10mer PNAs based on an *acpP*-targeting PNA. AcpP is an essential protein and its mRNA has been successfully targeted with PNAs and phosphorodiamidate morpholino oligomers (PMOs) before (Popella et al. 2021; Moustafa et al. 2021; Dryselius et al. 2003; Tilley et al. 2006). We introduced two adjacent mismatch positions from the N to the C terminus of the *acpP*-targeting PNA, while preserving the initial GC content of 40% (Fig. 2A, Supplemental Table S1). We refer to this series of mismatched PNAs by the position of the first mismatch, e.g., mm1 refers to the PNA with mismatches at positions one and two. Additionally we used a scrambled PNA to profile unspecific effects (Supplemental Table S1). All PNAs were fused to a KFFKFFKFFK peptide (KFF), a potent CPP for *Salmonella* (Popella et al. 2021).

**Figure 2.**
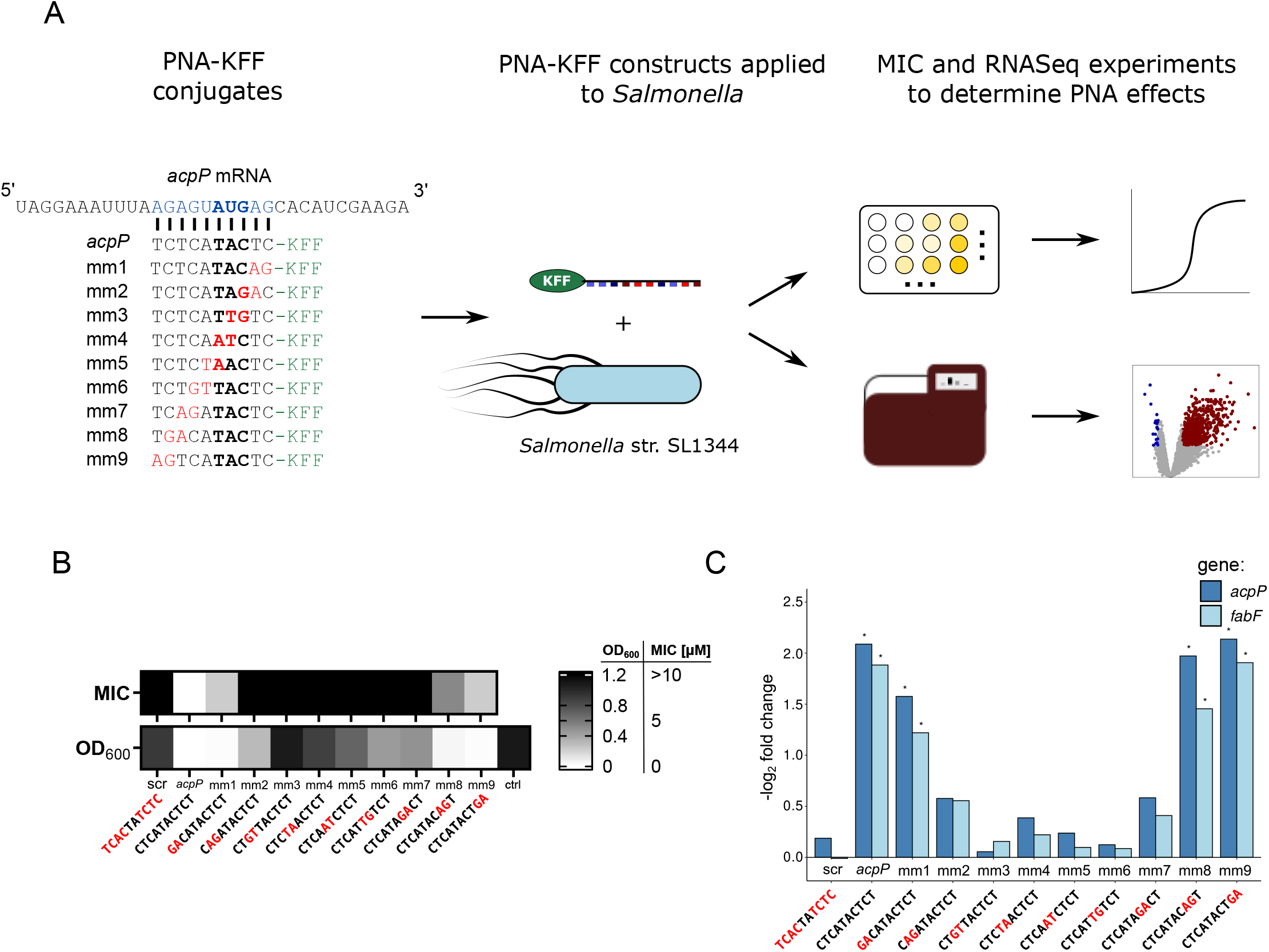
PNAs with two terminal mismatches show position-dependent effects on *Salmonella* growth and mRNA levels. **(A)** Workflow of experiments. *acpP* PNA (*acpP*) and nine *acpP*-related PNAs with two serial mismatches (mm1-mm9) were designed and tested against *S. enterica* serovar Typhimurium strain SL1344. PNA sequences are shown from their C-to N-terminus (from left to right). A 10-mer region of the *acpP* mRNA (blue) including its “AUG” start codon (bold) was targeted by one fully complementary PNA and nine mismatched PNAs, all coupled to the CPP KFF (green). Mismatches in the PNA sequences are marked in red. Upper right: Minimum inhibitory concentration experiments were performed by challenging *Salmonella* with different CPP-PNA concentrations for 24 hours while measuring cellular growth. Lower right: RNA sequencing was performed to profile RNA levels after short term CPP-PNA exposure. **(B)** Summary of growth experiments with *Salmonella* treated with KFF-PNAs. (Top) Heatmap of MIC values, determined as the lowest concentration that fully inhibits growth after 24 hours (see Figure S6). (Bottom) Heatmap of OD_600_ after 24 h treatment with 10 μM of the indicated PNA, or a water control. **(C)** Transcript levels of *acpP* and *fabF* after short-term exposure (15 minutes) to 5 μM of the indicated PNA relative to the untreated control condition. The barplot shows the negative log_2_ fold change of *acpP* (steel blue) and *fabF* (light blue) transcripts from the RNA-seq experiment. Asterisks indicate significant downregulation (log_2_FC > −1 & FDR <0.001).

First, we determined the minimum inhibitory concentration (MIC) of each PNA, defined as the lowest concentration that fully inhibits visible growth (OD_600 nm_ <0.1) of *Salmonella* (~10^5^ cfu/ml). Consistent with previous results (Popella et al. 2021), the PNA targeting the TIR of the *acpP* transcript with zero mismatches had an MIC of 1.25 μM. Two bp mismatches led to >4 fold increases in MIC values compared to the fully matched *acpP* PNA, most likely due to reduced binding affinity (Fig. 2B). Interestingly, we observed a mismatch-position dependent growth inhibition of *Salmonella* (Fig. 2B, Supplemental Fig. S6). PNAs that carry mismatches at either end of the sequence (i.e., mm1 and mm9) have MICs of 5 μM, whereas more central mismatch positions lead to MICs of 10 μM or higher (Fig. 2B). This is also seen in *Salmonella* growth curves (Supplemental Fig. S6), where growth defects become apparent at lower concentrations for mm1 and mm9 compared to PNAs with mismatches at more central positions. PNAs mm3 and mm4, which possess two mismatches to the start codon AUG, did not show any growth effect even at high concentrations, whereas mm1, mm8 and mm9 were still able to interfere with *Salmonella* growth (Supplemental Fig. S6).

### PNAs with terminal double-mismatches retain ability to trigger target mRNA depletion

Previously, we have shown that RNA-seq can serve as readout for direct effects on transcript levels after short-term treatment with a PNA targeting *acpP* in *Salmonella* (Popella et al. 2021). To analyze to what degree the designed series of PNAs affect on- and off-target mRNA levels compared to the “on-target” *acpP* PNA, we performed RNA-seq on *Salmonella* after a 15-minute exposure to 5 μM of the KFF-conjugated PNAs. To identify differentially expressed genes, we contrasted transcript levels in PNA-treated *Salmonella* with that of untreated control samples. First, we examined depletion of the target transcript, *acpP*, and *fabF*, a gene co-transcribed with *acpP* and shown to be concomitantly depleted by *acpP* PNA (Popella et al. 2021) (Fig. 2C). In contrast to the scrambled control, mRNAs of both *acpP* and *fabF* were significantly depleted by the fully-matching *acpP* PNA (log_2_FC < −1 and FDR < 0.001). mRNA depletion caused by mismatched PNAs correlated with the MIC values (Fig. 2B, C). In particular, PNAs mm1 and mm9, which both had an MIC of 5 μM, resulted in significant (log_2_FC < −1 and FDR <0.001) mRNA depletion of both *acpP* (log_2_FC = −1.6 (mm1) and −2.1 (mm9), FDR <0.001) and *fabF* (log_2_FC = −1.2 (mm1) and −1.9 (mm9), resp., FDR <0.001) (Fig. 2C). mm8 also induced significant depletion of both transcripts, indicating that only seven consecutively matching bases can be sufficient to affect target transcript levels. On the other hand, PNA sequences with central mismatches (mm3 through mm7) did not cause a significant depletion of the target mRNAs, consistent with their negligible effect on *Salmonella* growth (Fig. 2B, C). Altogether, these data show that PNAs with double mismatches in their termini and seven or more consecutive nucleobases available for base pairing retain antisense activity based on growth inhibition and depletion of *acpP* transcript levels.

### PNAs affect mRNA levels of off-target genes in a mismatch-dependent manner in both Salmonella and UPEC

Next, we analyzed the effects of our series of PNA variants on the complete *Salmonella* transcriptome. Principal component analysis (PCA) showed a clear separation on the first principal component between untreated control samples and the KFF-PNA conjugates, suggesting that the dominant effect in our data set was caused by KFF-PNA treatment itself, and not specifically due to depletion of *acpP* (Supplemental Fig. S7B). The global transcriptomic response to PNAs differs across PNA treatments and depends on the chosen PNA sequence (Fig. 3). Interestingly, more genes are upregulated than depleted in all samples (Fig. 3, Supplemental Fig. S8A). As shown previously in both *Salmonella* and UPEC, the most strongly upregulated genes after short-term KFF-PNA exposure are part of an envelope stress response (Supplemental Fig. S9, (Popella et al. 2021, 2022). In line with this observation, gene set analysis revealed that genes of the PhoPQ and PmrAB regulons as well as the KEGG-pathways “Two component system” and “cationic antimicrobial peptide (CAMP) response” are strongly induced upon PNA exposure for all PNA-conjugates (Supplemental Fig. S9).

**Figure 3.**
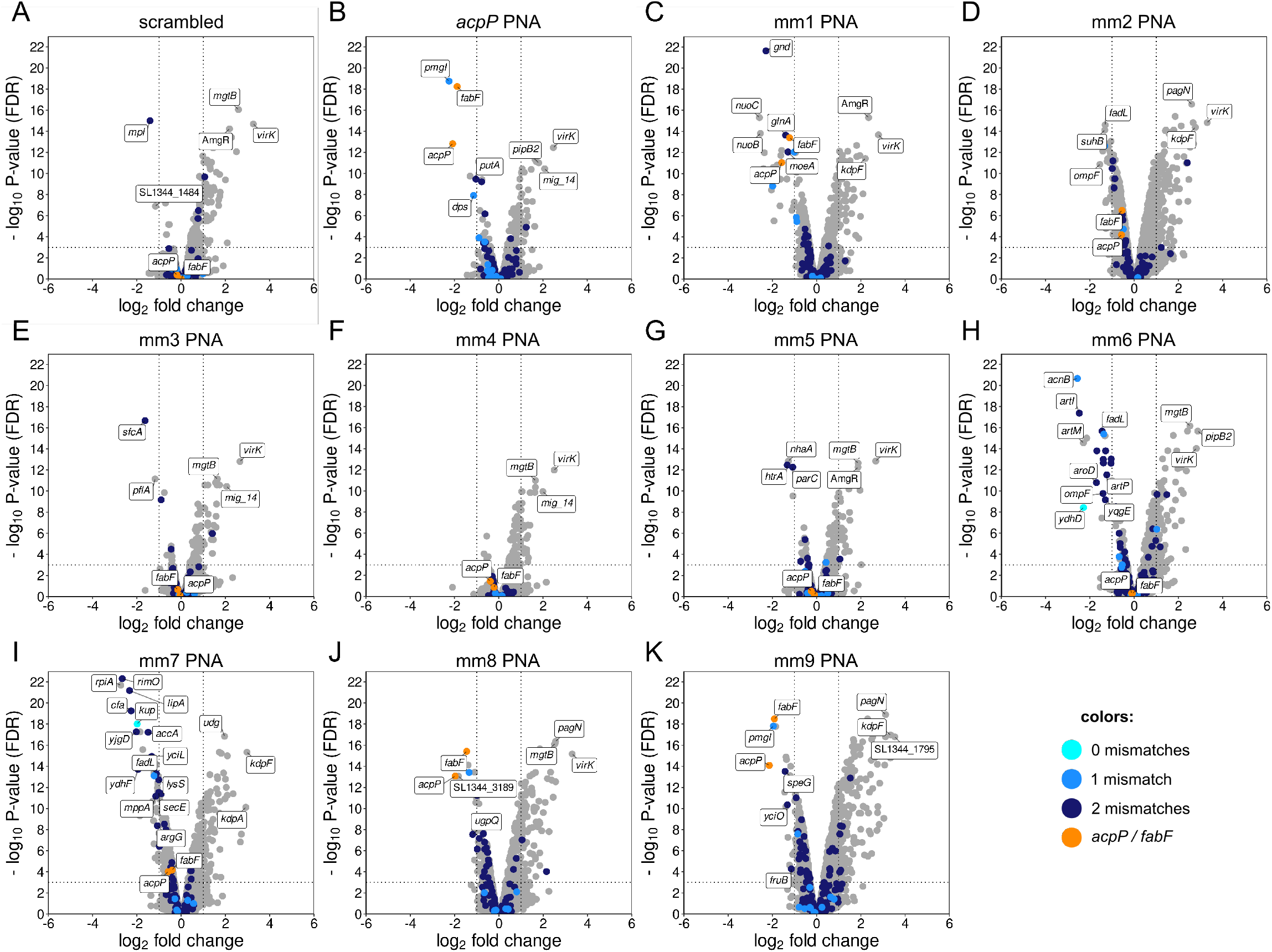
Transcriptomic profiles of *Salmonella* in response to 15 minutes of treatment by PNAs. Volcano plots show changes in *Salmonella* transcript levels as fold change (log_2_) compared to the untreated (water) control sample on the x-axis. The y-axis shows false discovery rate (FDR)-adjusted P-values (–log_10_, y-axis). A scrambled control PNA **(A)**, the fully matching *acpP* PNA **(B)** and all double-mismatch PNA samples **(C-K)** are shown. Significantly differentially expressed (DE) genes are characterized by an absolute fold change >2 (depleted if log_2_FC < –1, upregulated if log_2_FC > 1; vertical dashed line) and an FDR-adjusted P-value <0.001 (–log_10_FDR > 3, horizontal dashed line). Target genes (*acpP* and *fabF*) are labeled and highlighted in orange. Genes with zero-, one- and two-mismatch complementarity at the translation initiation region with the respective PNA are highlighted in cyan, blue and dark blue, respectively.

Contrary to the consistent pattern of upregulated genes, genes with significantly reduced mRNA levels differ between the PNA samples. This suggests that mRNA depletion might be PNA-sequence dependent (Fig. 4A). To determine whether the downregulation of transcripts is caused by off-target effects, we ran MASON to search for off-target sites for all PNAs in TIRs of genes and manually checked whether significantly depleted genes have complementarity with the respective PNA sequence. Interestingly, many significantly downregulated genes have complementarity to the PNA with up to 3 mismatches in their TIR (Fig. 3, Fig. 4A). In total, of 160 significantly depleted genes across all samples, we identified off-target sites for 72 (Supplemental Fig. S8B).

**Figure 4.**
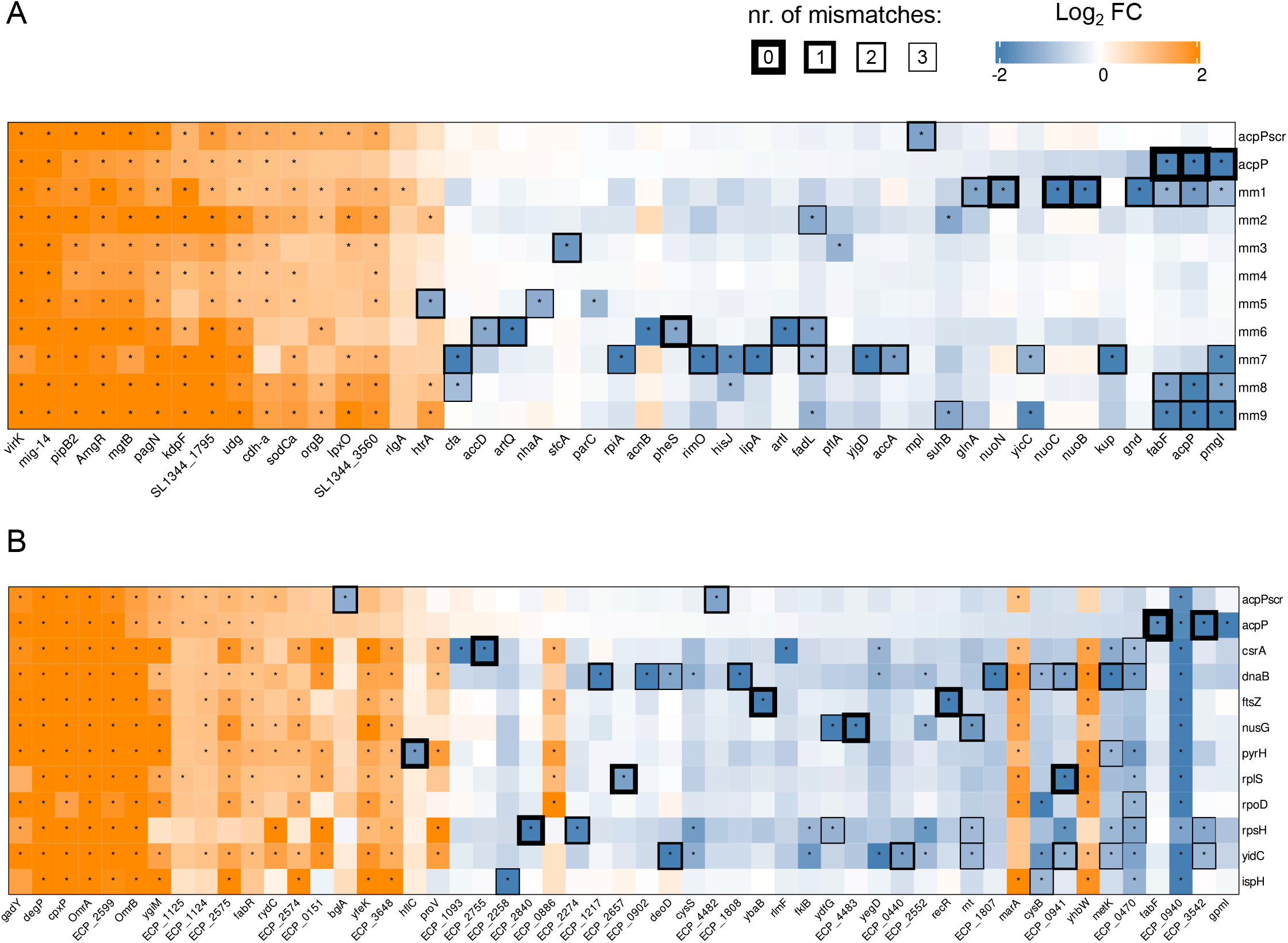
Changes in *Salmonella* **(A)** and UPEC **(B)** gene expression in response to PNA treatment, inferred from RNA-seq data. The heatmap shows genes which rank amongst the top 5 significantly enriched and depleted genes of each sample. Columns are ordered by decreasing log_2_FC in the sample treated with the fully matching *acpP* PNA. Colors indicate the log_2_ fold change of the respective gene after 15 minutes of PNA exposure, with orange and blue showing positive and negative differences in gene expression, respectively. Asterisks indicate significant changes (log_2_FC >1, FDR <0.001). Each row represents one PNA condition and each column a specific gene. The top five regulated genes (based on P-value) with an absolute fold change >2 and FDR <0.001 (marked with asterisks) are shown for each sample. Heatmap rectangles are framed if the PNA-gene pair has a predicted off-target with 0, 1, 2 or 3 mismatches. The frame thickness denotes the complementarity of the gene-PNA pair; the less mismatches the thicker the frame.

Based on *acpP* PNA-dependent depletion of *fabF*, the gene downstream of *acpP*, we also assigned off-target status to genes lying immediately downstream (<30 base gap) of an off-target gene. For these, we observed a similar co-regulation, for example, the mm1 PNA has complementarity with the *nuoA* TIR with one bp mismatch at position 10, and depletes the 12 genes downstream of the nuo operon as well as *nuoA* itself (Supplemental Fig. S10). These polar effects are likely caused by the transcription of multiple genes from a single promoter within bacterial operons.

To test whether these results are specific to *Salmonella*, we applied the same bioinformatics analysis to a published transcriptomic data set in which UPEC were treated with 11 different PNAs targeting various essential genes (Popella et al. 2022). The overall transcriptomic pattern closely resembles the data in the present study (Fig. 4A, B; Supplemental Fig. S8A, C). Specifically, we identified a number of consistently upregulated genes, many from outer membrane sensors and two component systems. This is likely due to the envelope stress response triggered by CPP-coupled PNAs (Fig. 4A, B, Supplemental Fig. S9, (Popella et al. 2021, 2022)). Of the 369 unique significantly depleted genes, we observed off-target sites in TIRs of 192 genes (Fig. 4B, Supplemental Fig. S8D). For example, the PNA targeting *ftsZ* had full complementarity to the TIR of *ybaB*, one of the significantly depleted genes. These comparisons suggest that, in both organisms, PNA complementarity to TIRs of non-target genes often leads to depletion of the respective mRNAs.

### Mismatch position influences the degree of off-target mRNA depletion

So far, we found that PNA complementarity with TIRs of off-target genes with few mismatches can induce mRNA depletion. We next wanted to test whether this effect depends on the specific position of the mismatches within the PNA sequence, as we saw with the PNA variants targeting *acpP* in *Salmonella*.

For predicted off-targets with 1-3 bp mismatches in *Salmonella* and UPEC, we determined the most central mismatch position. For instance, if there was an off-target binding site with mismatches at positions one and three, we counted the mismatch at position three. We then plotted the most central mismatch position against the percentage of off-targets that are significantly depleted in the RNA-seq datasets (Fig. 5A, D). In these plots, a value of one includes off-target genes with mismatches at positions one and/or 10 whereas a value of five includes all off-targets with their most central mismatch at positions four and/or five. Our results show that of all off-targets with mismatches at a terminal position 30% and 63% of genes are significantly depleted in *Salmonella* and UPEC, respectively. The fraction of significantly depleted genes decreases rapidly as the mismatch position moves closer to the center (Fig. 5A, D). These results suggest that a longer stretch of matching bases is more likely to trigger off-target mRNA-depletion, while central mismatches reduce the likelihood of an off-target effect.

**Figure 5.**
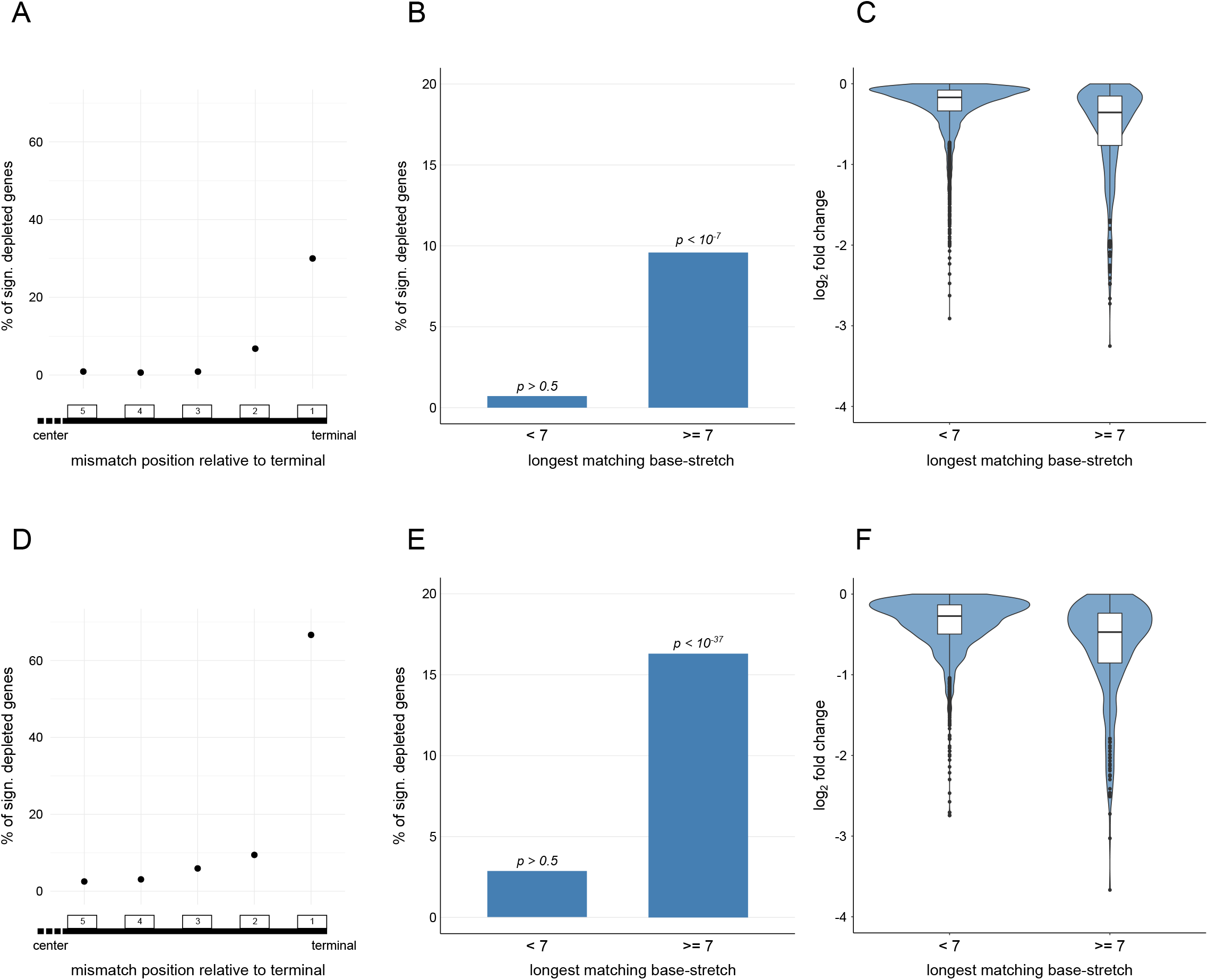
Mismatch position within the PNA target region dictates the degree of the off-target effect. The TIR of all genes were screened for complementarity to the PNA with a maximum of 3 mismatching bases. Genes with less than 5 counts per million were excluded from the analysis. For all off-targets with >0 mismatches, the most central mismatch was determined and plotted against the percentage of significantly downregulated genes for the *Salmonella* **(A)** and UPEC **(D)** data sets. Off-targets with a maximum base-pairing stretch (i.e. consecutive matches without mismatch) of >6 were compared to the rest of the off-targets in *Salmonella* **(B)** and UPEC **(E)**. Hypergeometric p-values on top of the bars test for enrichment of differentially depleted genes in the RNA-seq datasets relative to random draws with the same sample size. Log_2_ fold changes of off-target genes with <7 and >6 were additionally plotted for genes with log_2_FC les than 0 in *Salmonella* **(C)** and UPEC **(F)**.

To define a general rule for PNA design, we assessed whether the longest stretch of matching bases can be used as a predictor of off-target effects. We observed that for *Salmonella* and UPEC, 9.6% and 16.3% of all off-targets with at least seven consecutive matches were significantly depleted, respectively (Fig. 5B, E). This is likely an underestimate of the total number of effective off-targets, as we have recently shown that targeting of mRNAs by PNAs does not necessarily lead to depletion of the transcript (Popella et al. 2022). Low PNA-mRNA affinity appears to ameliorate off-target activity, as we found that requiring a predicted T_m_ of >20 °C increased the fraction of depleted off-targets to 13.6% and 25.1%, respectively (Supplemental Fig. S11). On the other hand, <5% of genes were depleted if they had an off-target site of six or less consecutive matches. Using a hypergeometric test we confirmed that the observed effects in the samples with at least seven consecutive matches cannot be explained by random draws (p <10^−7^ and p <10^−37^ for *Salmonella* and UPEC, resp.), whereas the <5% of depleted genes with off-target sites of six or less consecutive matches are consistent with a random draw (p >0.5) (Fig. 5B, E). We also observed a stronger overall depletion for transcripts with off-targets of at least seven consecutive matches (Fig. 5C, F). We therefore defined off-targets as “critical” only when they have at least seven consecutive matching bases and incorporated this finding into MASON (Fig. 1C). These data highlight that determining the number of consecutively matching bases can improve off-target prediction of PNAs in bacteria.

## DISCUSSION

Here we present MASON, a web-tool that facilitates the design of PNA sequences targeting bacterial genes. We also provide experimental evidence to justify MASON’s off-target prediction, which considers the mismatch position within the PNA. Combining growth measurements and RNA-seq analyses, we show that 10-mer PNAs targeting the essential gene *acpP* with terminal double mismatches retain their ability to inhibit *Salmonella* growth and to reduce transcript levels. Bioinformatic analysis of transcriptomic datasets reveal the effects of PNAs on transcripts with off-target binding sites and show how the position of mismatches impact these effects. Specifically, our analyses demonstrate that off-target binding sites can trigger mRNA depletion when at least seven consecutive bases match the PNA sequence. Our data argue for the importance of allowing terminal mismatches for off-target assessment of PNAs, especially when long stretches of the PNA can bind the mRNA.

There are various computational tools that aid the design of antisense oligonucleotides, RNAi, and CRISPR gRNAs that target eukaryotic genes (Chalk and Sonnhammer 2002; Sciabola et al. 2021; Owczarzy et al. 2008; Liu et al. 2020; Li et al. 2021). However, these methods are not suitable for designing short bacterial PNAs because these usually apply different design rules due to differences in binding affinity, mismatch tolerance, sequence length and mode of delivery. Tools for designing bacterial PNAs are rare. First, *PNA finder* is a toolbox that generates PNA sequences for a set of genes while considering issues like self-complementarity, solubility and off-target sites (Eller et al. 2021). A drawback of this method is that it only runs on a Bash shell, making it difficult to install for researchers without computational expertise. Second, *PNA Tool* is a freely accessible webserver which screens specified PNA sequences, predicts PNA-DNA T_m_ and analyzes other PNA-specific attributes like GC content and self-complementarity (https://www.pnabio.com/support/PNA_Tool.htm). However, it does not help in the design of sequences for a specific gene nor does it evaluate organism-specific target and off-target effects. By contrast, MASON is an easy to run webserver that designs bacterial PNA sequences for selected bacterial genes and predicts off-target effects as well as T_m_ and other attributes.

While we designed MASON for PNA sequences, it is likely similarly useful for the design of other ASOs, such as phosphorodiamidate morpholino oligomers (PMOs). PMOs are also nucleic acid mimics that only differ in their backbone compared to PNAs (Iversen 2001). Since PMOs take advantage of a similar mechanism for target gene inhibition (Deere et al. 2005; Geller et al. 2003), they could show similar off-target effects as well.

We showed that 10-mer PNAs with consecutive double mismatches are still able to inhibit the expression of *acpP*, if these mismatches are located at the termini of the PNA. We recently also observed this mismatch-position dependent off-target effect by cell-free *in vitro* ribosome profiling (Hör et al. 2022), adding further confidence to this observation. The fact that PNAs with as few as seven consecutive matches can retain activity suggests that the design of PNAs shorter than 10 nucleobases is feasible. Because PNA length is an important bottleneck that limits cellular entry (Goltermann et al. 2019), shorter PNAs might improve cellular uptake. While previous studies observed reduced PNA activity at a length of less than 10 bases (Good et al. 2001; Goltermann et al. 2019), we recently found that 9-mer PNAs targeting essential genes in UPEC were as efficient as their 10-mer counterparts in inhibiting bacterial growth in some cases (Popella et al. 2022).

The MASON webserver currently designs PNAs targeting mRNA sequences overlapping the TIR of a target gene, as this region has been previously shown to be most effective for PNA-based inhibition of translation. In a pioneering study in *E. coli*, Dryselius et al. designed PNAs targeting all positions in the mRNA of the *acpP* and *lacZ* genes, finding that only the regions around the SD sequence and the start codon are potent targets for inhibiting protein production (*lacZ*) or preventing cellular growth (*acpP*) (Dryselius et al. 2003). This position-dependent effect of PNAs led us to focus on off-targets in the TIR sequence, which could be particularly problematic when PNAs are used for gene silencing rather than killing target bacteria.

We will continue to apply data-driven methods to improve future versions of MASON in our effort to provide general design principles for bacterial ASOs, and we hope that MASON will become a widely used and evolving resource for the design of antisense drugs targeting bacterial pathogens.

## MATERIALS AND METHODS

### MASON web browser

MASON (Make Antisense Oligomers Now) is a webserver that provides researchers with ASO designs targeting bacterial genes of interest (Supplemental Fig. S1). There are five required user inputs (Supplemental Fig. S2). (i) A custom ID for later retrieval of the result; (ii) the genome and annotation of a bacterium of interest in fasta and gff format, or alternatively, selection of one of four pre-loaded bacteria comprising *Escherichia coli* str. K12 substr. MG1655, *Salmonella enterica* subsp. *enterica* serovar Typhimurium SL1344, *Clostridium difficile* 630 and *Fusobacterium nucleatum* ATCC 23726; (iii) the locus tag of one or more target genes, or for pre-loaded organisms, a selection from a dropdown menu of experimentally determined essential genes; (iv) the length of the designed ASOs; and (v) the allowed number of mismatches for off-target screening. In default settings, only ASOs overlapping the start codon are designed. Optionally, additional 5’ UTR sequences upstream of the start codon can be selected to target the RBS, and genes from the human genome or the microbiome can be additionally screened for off-targets.

MASON then runs for 10-120 seconds per target gene before returning the output. The output consists of three main elements (Supplemental Fig. S3). First, a heatmap visualizing all designed ASO sequences is shown, where the target mRNA and alignment of each ASO is visualized along with the respective start codon (Fig. 1B, Supplemental Fig. S3A). All possible ASOs spanning the target region are designed, with the exception of ASOs with more than 60% self-complementarity. Self-complementarity was defined as the maximal number of consecutive Watson and Crick matches of the ASO with itself, aligned in either direction. Secondly for each ASO sequence, T_m_ between mRNAs and ASOs are visualized in a barplot (Supplemental Fig. S3B). Third, the number of predicted off-target sites for each ASO are visualized in a barplot, summarizing off-targets in both the whole transcriptome, as well as in the TIR of other genes in the selected organisms (Fig. 1C, Fig. S3C). Finally, an additional summary table reports all aforementioned attributes along with general features such as position from start site and self-complementarity (Supplemental Fig. S5). The web interface was written using the python-based web-framework Flask (Grinberg 2018). The MASON website can be freely accessed at https://www.helmholtz-hiri.de/en/datasets/mason. The code can be viewed and accessed freely at https://github.com/BarquistLab/mason.

### MASON algorithm

The code was written in Python (v3.7), R (v4.1.1) with various modules from the Biopython and Bioconductor environments (Cock et al. 2009; Huber et al. 2015). Briefly, MASON takes the target gene and designs all possible ASO sequences which are complementary to the region overlapping the start codon of the target gene and the specified upstream region. Next, the number of self-complementary consecutive bases are calculated with the SequenceMatcher package from the difflib python package. If the length of self-complementarity accounts for more than 50% of the ASO-length the respective sequence is dropped and not used for further analysis. T_m_ of the ASO-target mRNA duplex are calculated using the R package rmelting (v1.8.0) of the MELTING 5 platform (Dumousseau et al. 2012). Specifically, the melting function is applied for calculating the T_m_ of RNA-RNA duplexes using a nearest neighbor algorithm (Xia et al. 1998).

For identifying off-target sites in the transcriptome, the supplied genome files are screened for motifs similar to the target region of the respective ASO. Specifically, the SeqMap tool was used to map the targeted mRNA sequence to mRNA sequences, with the specified option for allowed mismatches (Jiang and Wong 2008). The allowed mismatches are chosen in the user input in the start form of the webserver and are added to the Seqmap command (Supplemental Fig. S2). Two to three mismatches were used here and were shown to be a good option to capture off-targets for 10-mer PNAs. Only “critical” off-targets with at least 7 consecutive matching bases are kept for the analysis. For screening total off-targets, the whole transcript sequence of each annotated gene is used. Off-target sites in the TIR of transcripts are defined as binding regions with the first base in the region between −20 and +5 bases relative to the annotated start site of the CDS.

The off-target screen for the human microbiome was created with data from the HMP (Human Microbiome Project Consortium 2012). The genome and annotation files of all available complete genomes (2,179) were downloaded from the NCBI BioProject collection. Next, start regions of all annotated transcripts (in total 5,515,510 regions) were extracted and used for the additional off-target screen, using custom Linux bash commands. The human genome (GRCh38) was downloaded from the NCBI and full transcript sequences were extracted using the getfasta command of the BEDTools suite (Quinlan and Hall 2010). Finally, the results are visualized as described in the previous section.

### MASON command line tool

The command line version of MASON can be accessed and downloaded from https://github.com/BarquistLab/mason_commandline. It can be run from any Linux environment after installing all dependencies. The tool uses the same algorithms/code as the web tool with two additions: (i) the user has the option to specify an ASO sequence without any input gene. This option prevents MASON from designing new ASO sequences and instead uses the input ASO sequences for off-target analysis; (ii) there is an option to specify additional non-target organisms which are screened for off-targets.

### Bacterial strains and peptide nucleic acids (PNAs)

*Salmonella enterica* serovar Typhimurium strain SL1344 (provided by D. Bumann, MPIIB Berlin, Germany hoi (Hoiseth and Stocker 1981); internal strain number JVS-1574) was used throughout this study. *S. enterica* SL1344 was cultured in non-cation adjusted Mueller-Hinton Broth (MHB, BD Difco^™^, Thermo Fisher Scientific) with aeration at 37 °C and 220 rpm shaking. PNAs conjugated to the peptide KFFKFFKFFK (KFF) were obtained from Peps4LS GmbH. Quality and purity control of these constructs was performed by mass spectrometry and HPLC (purity >98 %). PNAs (Supplemental Table S1) were dissolved in nuclease-free ultrapure water and heated at 55 °C for 5 min prior to use. By measuring the absorbance at 260 nm using a NanoDrop spectrophotometer (A _260 nm_; ThermoFisher) the concentration of PNA solutions were calculated based on their extinction coefficient. Low retention pipette tips and low binding Eppendorf tubes (Sarstedt) were used for all PNA solutions. PNAs were stored at − 20 °C.

### PNA selection and design

A 10-mer PNA targeting the start-codon (−5 to +5 relative to CDS start) of the *acpP* gene served as a positive control. Nine additional PNAs were designed, containing the same sequence but with double mismatches starting in each possible location from one to nine (Fig. 2A, Supplemental Table S1). The mismatch PNAs were designed so that the GC-content was the same as the control sequence, at 40%. That is, for each mismatch, guanines were swapped to cytosines and adenines to thymines, and vice versa (Fig. 2A, Supplemental Table S1). The number of off-targets were analyzed using the MASON command line tool.

### Minimum inhibitory concentration (MIC) determination

For MIC determination the broth microdilution method was applied according to the Clinical and Laboratory Standards Institute guidelines and slightly modified from a recently published protocol (Goltermann and Nielsen 2020). In brief, an overnight culture of bacterial cells was diluted 1:100 in fresh MHB and grown to OD_600_ 0.5. After diluting the culture to approximately 10^5^ cfu/ml in MHB (1:2000-1:2500), 190 μl were dispensed into a 96-well plate (Thermo Fisher Scientific). Subsequently, 10 μl of a 20x PNA working solution was added to the respective well to adjust final concentrations from 10-0.3 μM. As a growth control, 10 μL of sterile nuclease-free water were added to the bacterial suspension. Growth was monitored over 24 hours by measuring the optical density (OD) at 600 nm every 20 min in a Synergy H1 plate reader (Biotek) with continuous double-orbital shaking (237 cpm) at 37 °C. The MIC of a PNA was determined as the lowest concentration that inhibited visible growth in the wells (OD (600 nm) <0.1).

### PNA treatment of S. enterica SL1344 for isolation of total RNA

Overnight cultures of *S. enterica* SL1344 were diluted 1:100 in fresh MHB and grown to an OD_600_ of 0.5. Afterwards, cultures were diluted to approximately 10^6^ cfu/ml in fresh MHB (1:100). Subsequently, 1.9 mL of the bacterial solution were transferred into 5 mL low-binding tubes (LABsolute) and 100 μL of a 20x PNA solution was added to reach a final concentration of 5 μM for each tested KFF-conjugated PNA. In parallel, an equal volume of sterile nuclease-free water, which was used as solvent for the test compounds, was added to the bacterial suspension and served as negative control. After a 15-min incubation at 37 °C, RNAprotect Bacteria (Qiagen) was added to the samples according to the manufacturer’s instructions. After 10 min, cells were pelleted at 21,100 *g* and 4 °C for 20 min. Pellets were either directly subjected to RNA isolation or stored at −20 °C (<1 day) for subsequent processing.

Total RNA was isolated from bacterial cell pellets using the miRNeasy Mini kit (Qiagen) according to protocol #3 previously described in Popella *et al*. (Popella et al. 2021). In brief, pellets were resuspended in TE buffer (pH 8.0) supplemented with 0.5 mg/mL lysozyme (Roth) and incubated for 5 min with repeated vortexing in between. After adding β-mercaptoethanol-containing RLT buffer and ethanol according to the manufacturer’s instructions, samples were loaded on the columns. Wash-steps were performed according to the manual. RNA concentration was measured using a NanoDrop spectrophotometer.

### RNA-sequencing (RNA-seq)

RNA samples were delivered to Core Unit SysMed (Julius-Maximilian-University Würzburg, Germany) for RNA-seq. Briefly, after treating RNA samples with DNase, sufficient RNA quality was verified on a bioanalyzer (RNA chip Agilent). Ribosomal RNA was depleted (RiboCop-META kit, Lexogen) and RNA was then reverse transcribed for cDNA library preparation using the Corall kit (Lexogen) according to the manufacturer’s instructions. After pooling cDNA library samples at equimolar amounts, quality was verified using a bioanalyzer (DNA chip Agilent). The cDNA pools were sequenced using the NextSeq 500 system (HighOutput flow cell, 400 M, 1x 75 cycle single-end; Illumina).

### RNA-seq read quantification

Read mapping and differential expression analysis were performed as previously described (Popella et al. 2021). Briefly, reads obtained from RNA sequencing were trimmed, filtered and mapped against the respective reference genome. The reference genome consisted of the the *Salmonella enterica* subsp. *enterica* serovar Typhimurium SL1344 (FQ312003.1) reference genome and three plasmids: pSLT_SL1344 (HE654724.1), pCol1B9_SL1344 (HE654725.1), and pRSF1010_SL1344 (HE654726.1) (Kröger et al. 2012). Bases with a Phred quality score of <10 were trimmed and adapters were removed using BBDuk. Next, the trimmed reads were mapped against the reference genome using BBMap (v38.84) and then assigned to genomic features, including both CDSs and annotated sRNAs (Kröger et al. 2012; Hör et al. 2020) using the featureCounts method of the Subread (2.0.1) package (Liao et al. 2014). Read mapping and quantification for the UPEC data was performed similarly, as described in (Popella et al. 2022).

### Differential expression analysis

Downstream analysis was performed using R (v4.1.2) and various packages from the Bioconductor project. Raw read counts were imported and analyzed with edgeR (v3.34.1) (Robinson et al. 2010). Genes with a cutoff value less than 10/L in at least three libraries were filtered, where L is the minimum library size in millions of counts, as proposed in (Chen et al. 2016). Raw read counts were normalized with the trimmed mean of M values (TMM) normalization (Robinson and Oshlack 2010). Filtered libraries were then examined for batch effects as proposed in (Peixoto et al. 2015). A consistent batch effect was identified between the two batches, which was corrected for by including a batch variable in the design matrix (Supplemental Fig. S7). Differential expression analysis was conducted using edgeR (Robinson et al. 2010). Estimation of quasi-likelihood (QL) dispersions was performed with the glmQLFit function. Then contrasts between all samples versus the water controls were created as input for the glmQLTest function. Genes with an absolute fold change > 2 and an adjusted P-value (Benjamini–Hochberg, (Benjamini and Hochberg 1995)) <0.001 were considered differentially expressed. The differentially expressed genes were visualized by volcano plots and heatmaps created using ggplot2 (v3.3.0) and the ComplexHeatmap (v2.4.2) package, respectively. The differential expression analysis for the UPEC data was performed in a similar way (Popella et al. 2022).

### Mismatch analysis

For the results shown in Figure 4, a modified analysis of off-target genes was performed. Each PNA sequence was screened for off-target matches using the MASON command line tool, while accepting all off-targets with 0-3 mismatches. Then, all significantly downregulated genes were framed if they had an off-target site in the TIR of the respective gene, or in a TIR of a gene located immediately lying immediately upstream (<30 base gap) of the gene. The frame thickness indicates that 0, 1, 2 or 3 mismatches were present in the off-target site.

### KEGG pathway enrichment analysis

To identify KEGG pathways for each gene, the R package KEGGREST (v1.28.0) was used. Additionally, gene sets of regulons curated in (Westermann et al. 2016) were added prior to the analysis. Rotation gene set testing (fry version of the roast gene set enrichment test (Wu et al. 2010)) was performed to identify enrichment of gene sets. Gene sets containing >10 genes with FDR-corrected P-values <0.001 are shown in Supplemental Figure S9, together with the median log_2_FC of genes in the respective pathway. If a sample had >10 significantly enriched gene sets, only the 10 gene sets with the lowest FDR adjusted P-values are shown.

### Off-target analysis of melting temperature, RNA abundance and RNA secondary structure

For Supplemental Figure S11, log_2_ fold changes and FDR-corrected p-values of transcripts with off-targets with at least seven consecutive matching bases were considered. Log_2_ fold changes and average log counts per million were calculated by edgeR, as described above. The T_m_ for off-target matches was calculated with the MELTING package as described before for the longest matching stretch in the of-target binding site (Dumousseau et al. 2012). To investigate the secondary structure of mRNAs surrounding the translation start site (−30 to +15 nucleotides relative to the CDS start), the minimum free energy (MFE) was calculated using the RNAfold command of the ViennaRNA software suite (Lorenz et al. 2011).

## DATA AVAILABILITY

The code of both the web- and the command line version MASON are published under the MIT License (v1.0) on Github https://github.com/BarquistLab/mason and https://github.com/BarquistLab/mason_commandline, respectively. The RNA-seq datasets from this study are available in GEO with the accession numbers GSE199542 and GSE191313. The up- and downstream analysis of RNA-seq data is available on Github https://github.com/BarquistLab/mason_paper.

## SUPPLEMENTAL MATERIAL

Supplemental material is available for this article.

## ACKNOWLEDGEMENTS

We thank Barbara Plaschke for technical assistance and Tobias Kerrinnes for laboratory organization. We thank Kristina Popova for contributions in the early phase of the project. We especially thank Anke Sparmann for useful comments and edits on the manuscript. We further thank Michael Kütt for helping with adding the webserver to the internet and providing useful comments on web development. We also thank the Core Unit Systems Medicine (Julius-Maximilians-University, Würzburg) for handling the RNA samples for sequencing.

