## Supplemental material for "Design and off-target prediction for antisense oligomers targeting bacterial mRNAs with the MASON webserver"

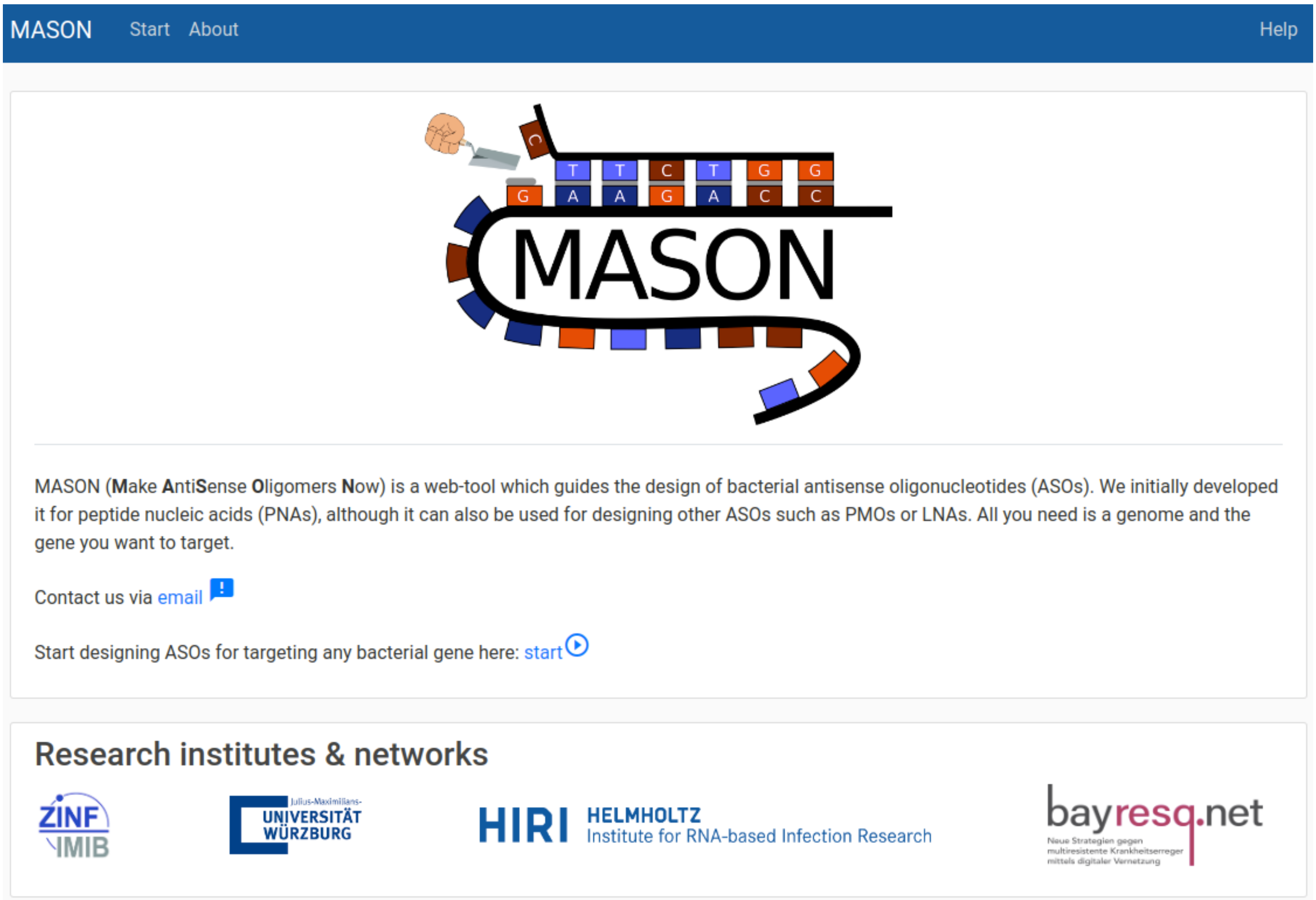


**Figure S1.** Web-interface of MASON showing the start homepage of MASON.

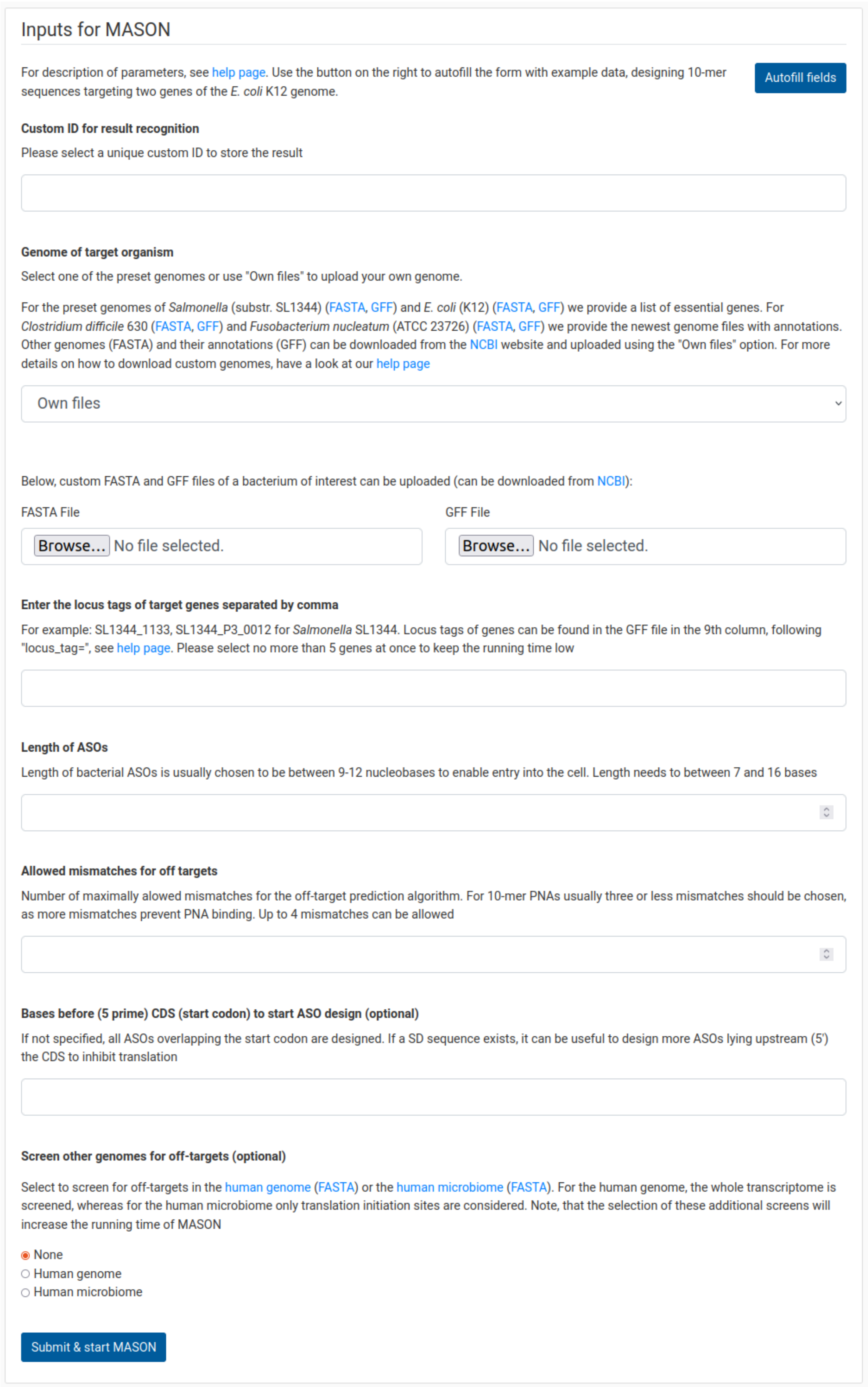
### Inputs for MASON

For description of parameters, see help page. Use the button on the right to autofill the form with example data, designing 10-mer sequences targeting two genes of the *E. coli* K12 genome.

Autofill fields

**Custom ID for result recognition**

Please select a unique custom ID to store the result

**Genome of target organism**

Select one of the preset genomes or use "Own files" to upload your own genome.

For the preset genomes of *Salmonella* (substr. SL1344) (FASTA, GFF) and *E. coli* (K12) (FASTA, GFF) we provide a list of essential genes. For *Clostridium difficile* 630 (FASTA, GFF) and *Fusobacterium nucleatum* (ATCC 23726) (FASTA, GFF) we provide the newest genome files with annotations. Other genomes (FASTA) and their annotations (GFF) can be downloaded from the NCBI website and uploaded using the "Own files" option. For more details on how to download custom genomes, have a look at our help page

Own files

Below, custom FASTA and GFF files of a bacterium of interest can be uploaded (can be downloaded from NCBI):

FASTA File

Browse... No file selected.

GFF File

Browse... No file selected.

**Enter the locus tags of target genes separated by comma**

For example: SL1344_1133, SL1344_P3_0012 for *Salmonella* SL1344. Locus tags of genes can be found in the GFF file in the 9th column, following "locus_tag=", see help page. Please select no more than 5 genes at once to keep the running time low

**Length of ASOs**

Length of bacterial ASOs is usually chosen to be between 9-12 nucleobases to enable entry into the cell. Length needs to between 7 and 16 bases

**Allowed mismatches for off targets**

Number of maximally allowed mismatches for the off-target prediction algorithm. For 10-mer PNAs usually three or less mismatches should be chosen, as more mismatches prevent PNA binding. Up to 4 mismatches can be allowed

**Bases before (5 prime) CDS (start codon) to start ASO design (optional)**

If not specified, all ASOs overlapping the start codon are designed. If a SD sequence exists, it can be useful to design more ASOs lying upstream (5') the CDS to inhibit translation

**Screen other genomes for off-targets (optional)**

Select to screen for off-targets in the human genome (FASTA) or the human microbiome (FASTA). For the human genome, the whole transcriptome is screened, whereas for the human microbiome only translation initiation sites are considered. Note, that the selection of these additional screens will increase the running time of MASON

- None
- Human genome
- Human microbiome

Submit & start MASON

**Figure S2.** User input page of the MASON webserver. All non-optional fields must be filled to start the calculation process of MASON.

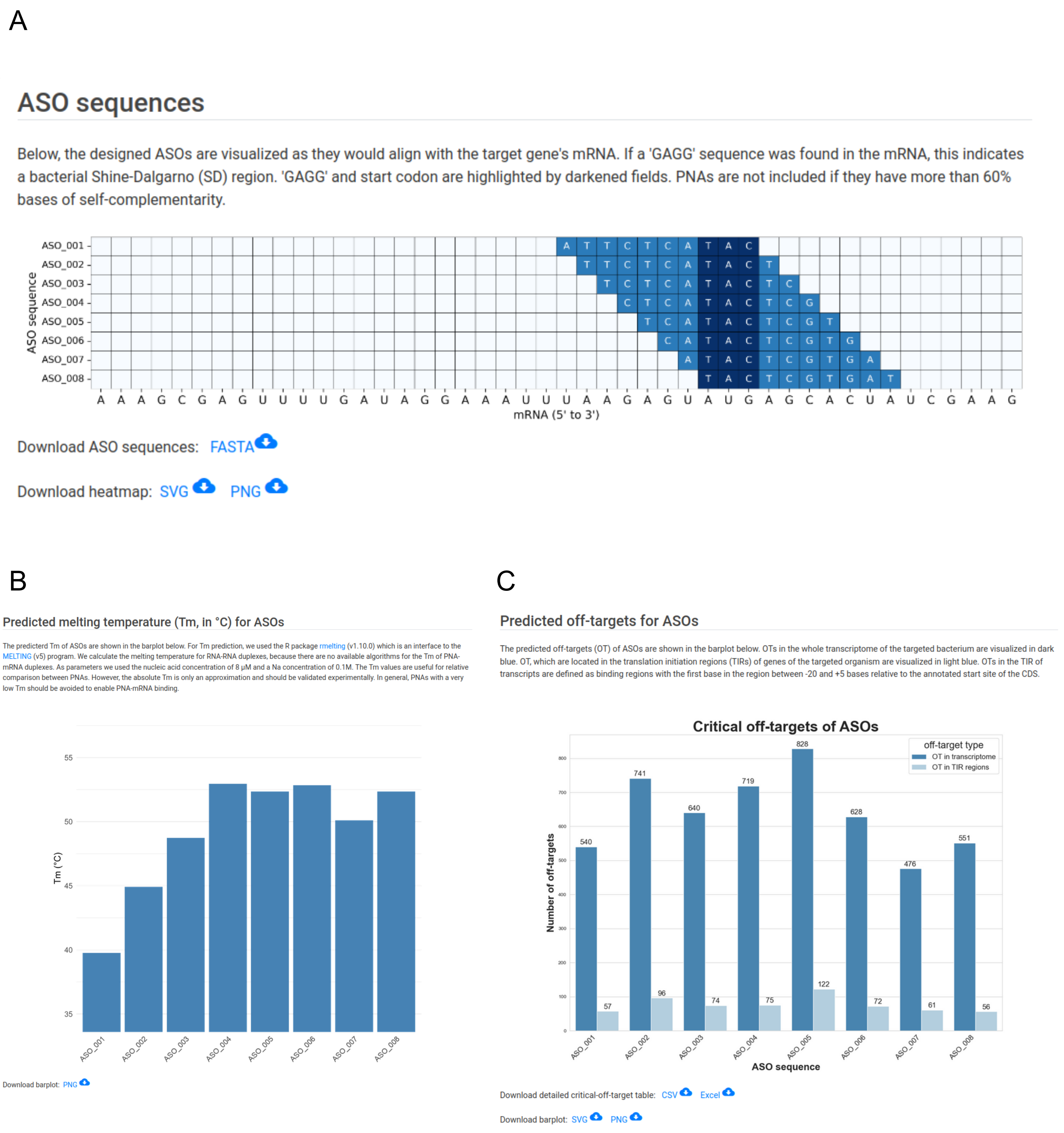


**Figure S3.** MASON results page. (A) Designed sequences targeting the AUG start codon. (B) Predicted PNA/mRNA melting temperatures of the designed sequences. (C) MASON's off-target predictions, showing critical off-targets in translation initiation regions and the whole transcriptome.

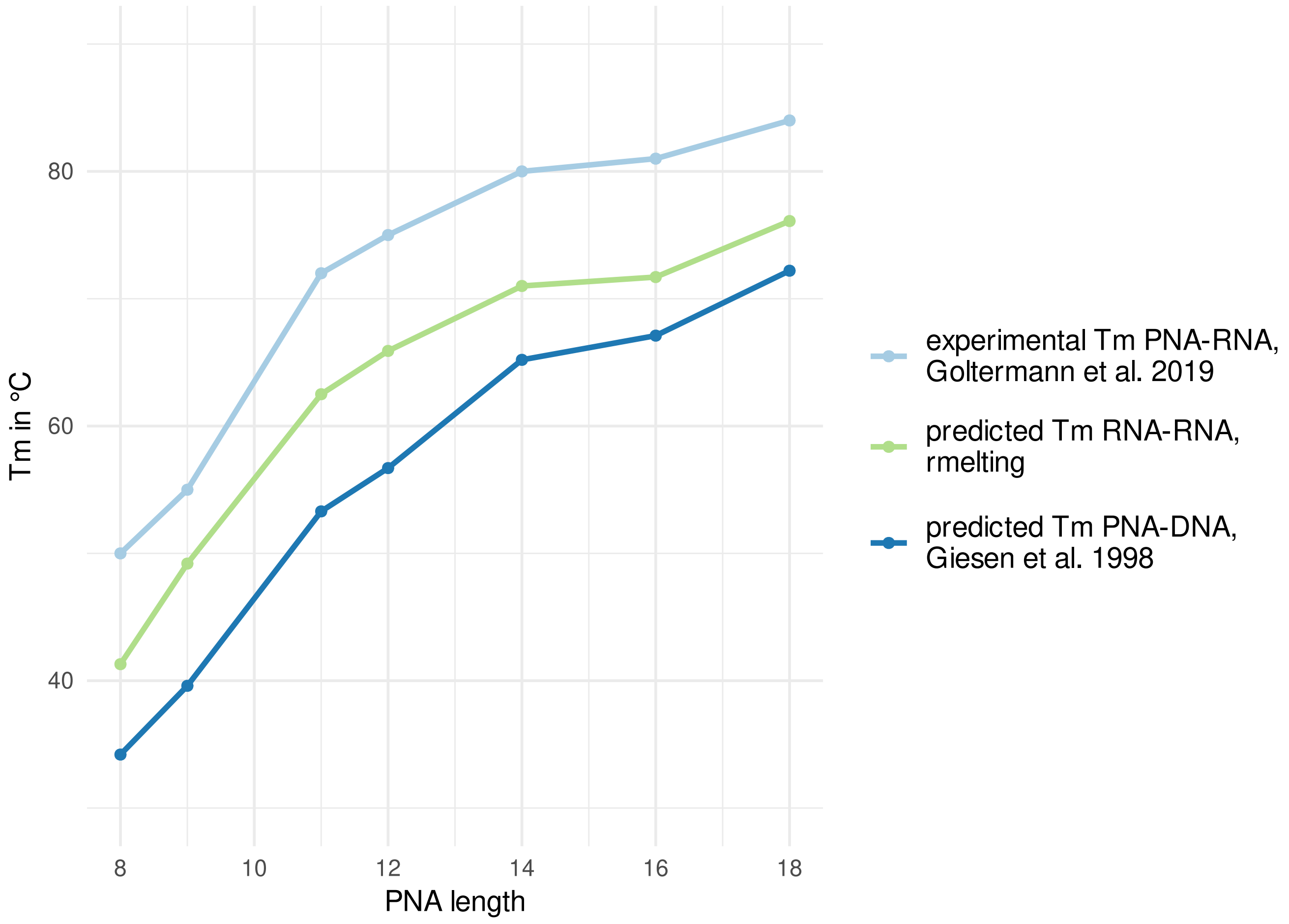


**Figure S4.** Comparison of predicted melting temperatures of rmelting (used by MASON) and the function of Giesen et al. (1998) compared to experimentally determined PNA-RNA duplex data from Goltermann et al. (2019).

### Summary table for - ASOs

Summary for each ASO sequence. meanings of column names:
location = location (start and end) of the ASO-binding site, measured from the CDS start site
SC_bases = maximum stretch of self-complementary bases
pur_perc = percentage of purine bases in ASO sequence (too high % should be avoided)
long_pur_stretch = longest purine stretch (too long purine stretches should be avoided)
Tm = Melting temperature
OT_tot = critical off-targets in the whole transcriptome
OT_TIR = critical off-targets in translation initiation regions

| ASO | ASO_seq | target_seq | location | SC_bases | pur_perc | long_pur_stretch | Tm | OT_tot | OT_TIR |
|---|---|---|---|---|---|---|---|---|---|
| ASO_001 | CATACTCTTA | UAAGAGUAUG | -7;3 | 2 | 30 | 1 | 39.76 | 540 | 57 |
| ASO_002 | TCATACTCTT | AAGAGUAUGA | -6;4 | 2 | 20 | 1 | 44.93 | 741 | 96 |
| ASO_003 | CTCATACTCT | AGAGUAUGAG | -5;5 | 2 | 20 | 1 | 48.73 | 640 | 74 |
| ASO_004 | GCTCATACTC | GAGUAUGAGC | -4;6 | 2 | 30 | 1 | 52.96 | 719 | 75 |
| ASO_005 | TGCTCATACT | AGUAUGAGCA | -3;7 | 2 | 30 | 1 | 52.36 | 828 | 122 |
| ASO_006 | GTGCTCATAC | GUAUGAGCAC | -2;8 | 2 | 40 | 1 | 52.84 | 628 | 72 |
| ASO_007 | AGTGCTCATA | UAUGAGCACU | -1;9 | 3 | 50 | 2 | 50.1 | 476 | 61 |
| ASO_008 | TAGTGCTCAT | AUGAGCACUA | 0;10 | 3 | 40 | 2 | 52.36 | 551 | 56 |

**Figure S5.** MASON results output table. This table summarizes sequence properties and off-target predictions for the designed ASOs.

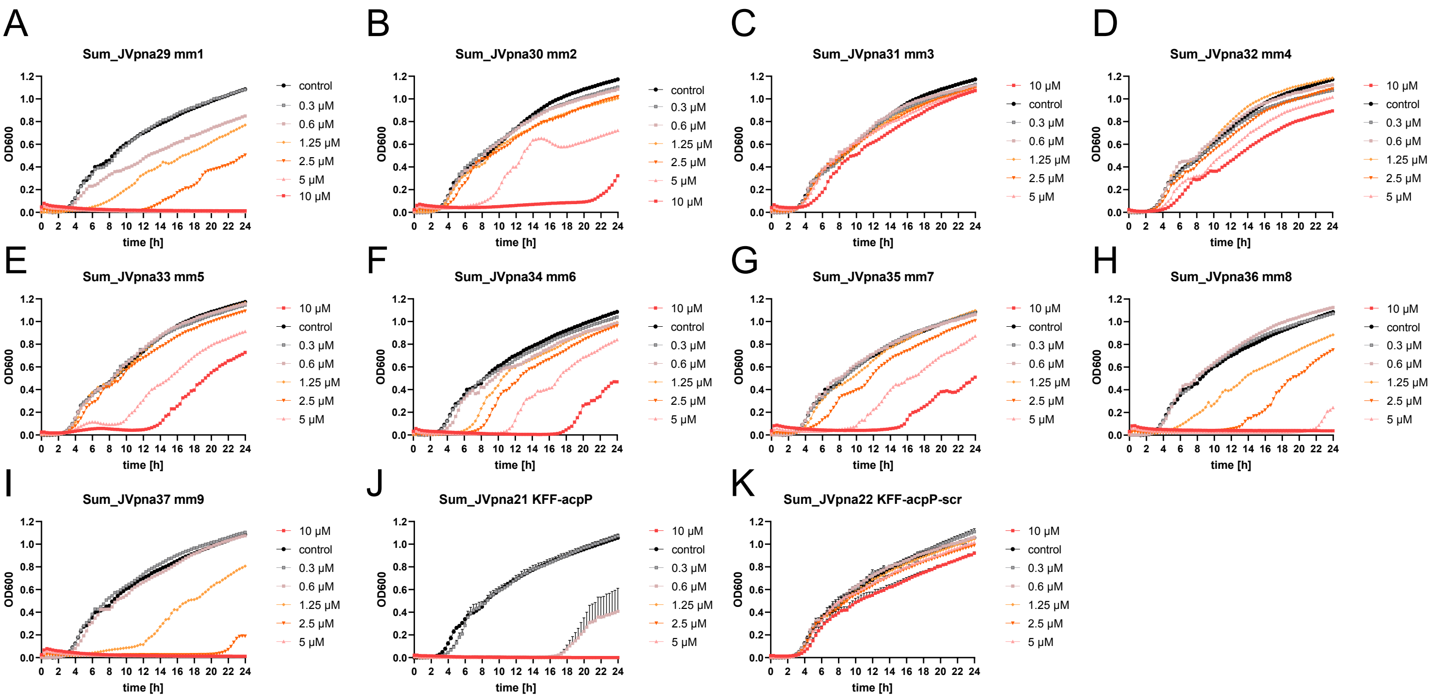


**Figure S6.** Effect of *acpP* mutagenesis PNAs with 2 base pair mismatch PNAs on *Salmonella enterica serovar* Typhimurium growth kinetics. Growth curves were measured as $OD_{600}$ over time using ~$10^6$ cfu/ml Salmonella while varying the concentrations of PNA conjugates. The experiments were performed in triplicate and error bars depict the standard error of the mean.

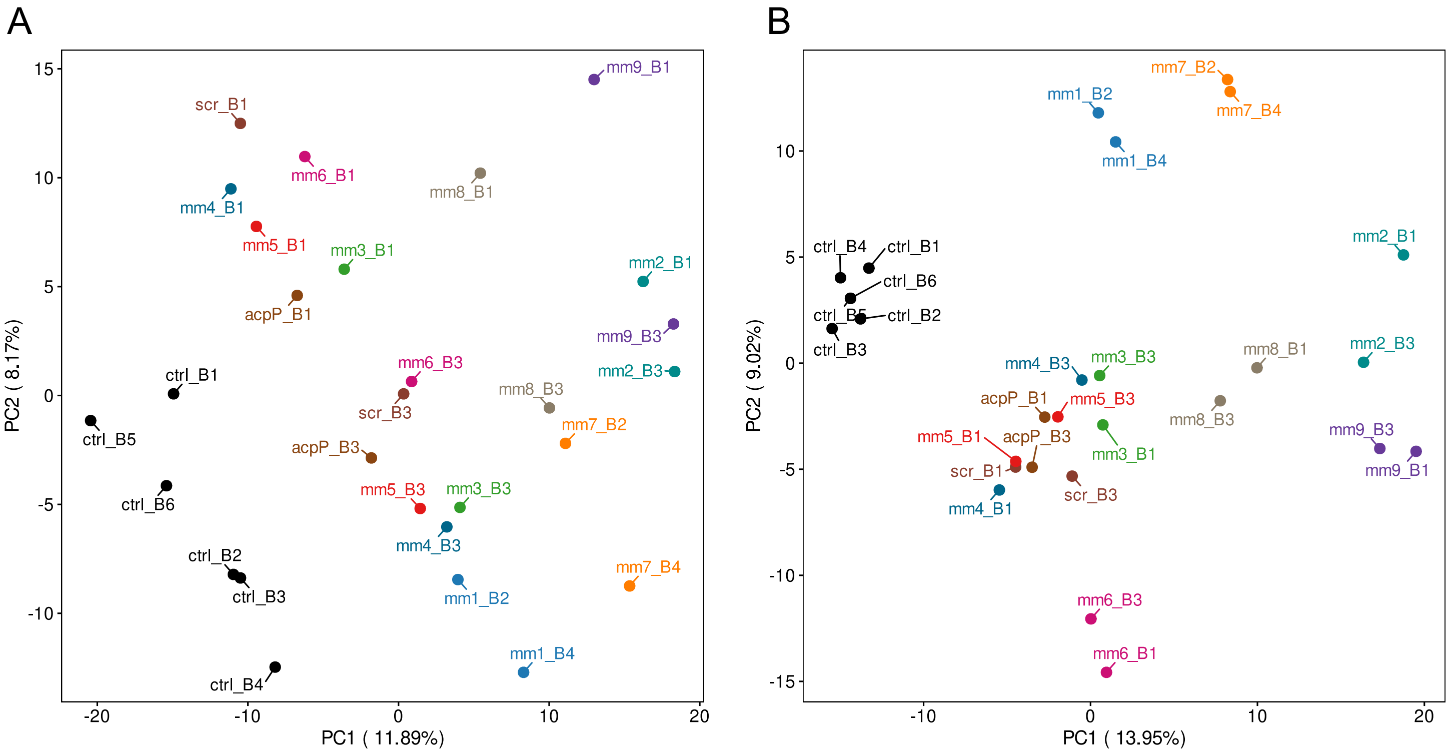


**Figure S7.** Principal component analysis (PCA) plots of the  of all 12 conditions of the *acpP* scanning mutagenesis experiment pre- and post- batch effect removal. Each of the CPP-PNA treated samples have two replicates, the untreated controls have 5 replicates. (A) PCA plot of TMM normalized read counts without batch-effect removal. (B) PCA plot after batch-effect removal. Batches are denoted with the batch number after B (B1-B6); colours mark treatment group.

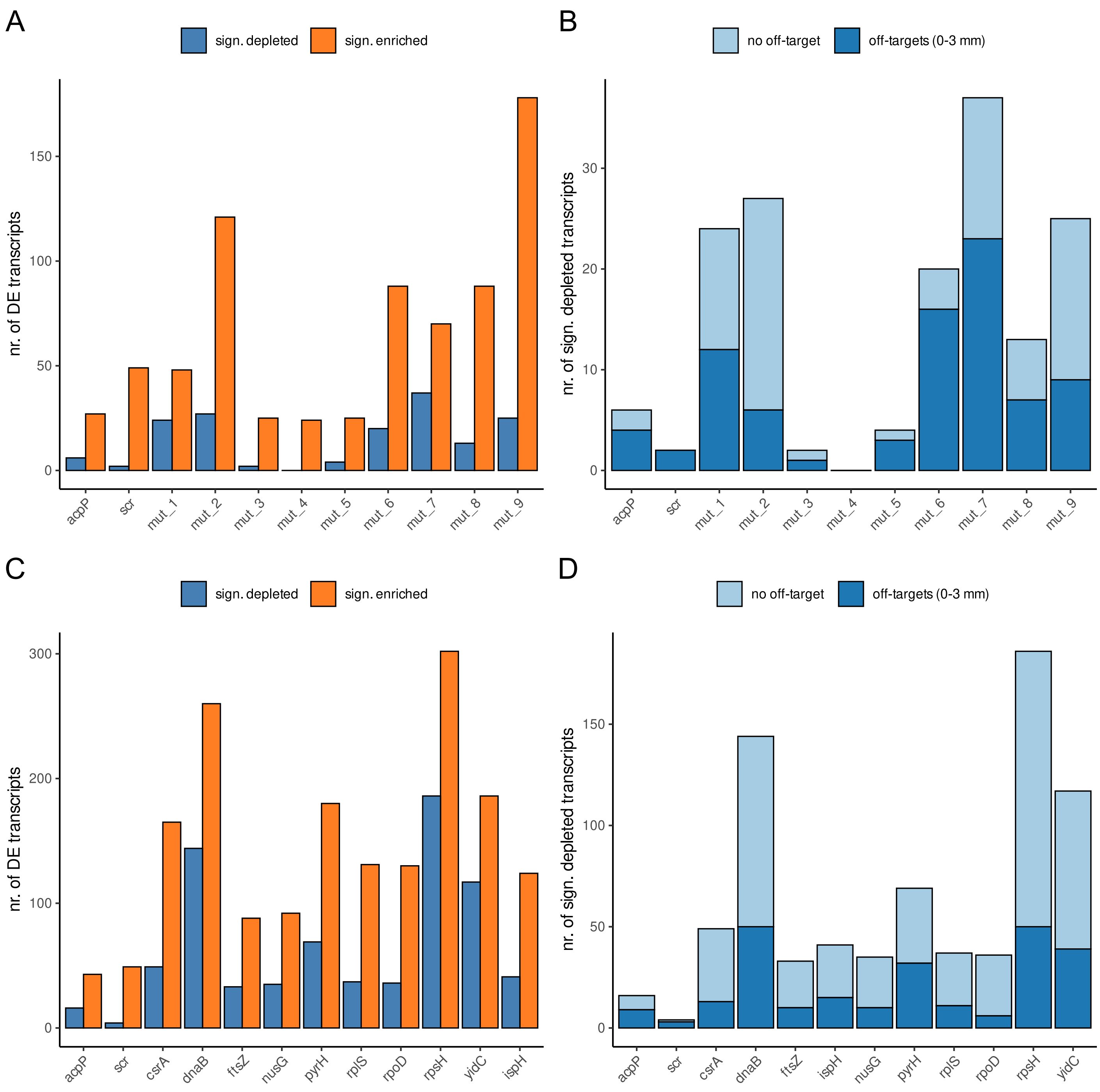


**Figure S8.** Number of up- and downregulated DE genes and ratio of off-targets in significantly depleted genes. (A, C) Number of differentially enriched (orange) and depleted (blue) transcripts for all transcriptomic experiments for *Salmonella* (A) and UPEC (C). Of the significantly depleted genes (blue bars), the amount of transcripts with off-target sites with up to three mismatches is plotted (darkblue) in a barplot for *Salmonella* (B) and UPEC (D) data.

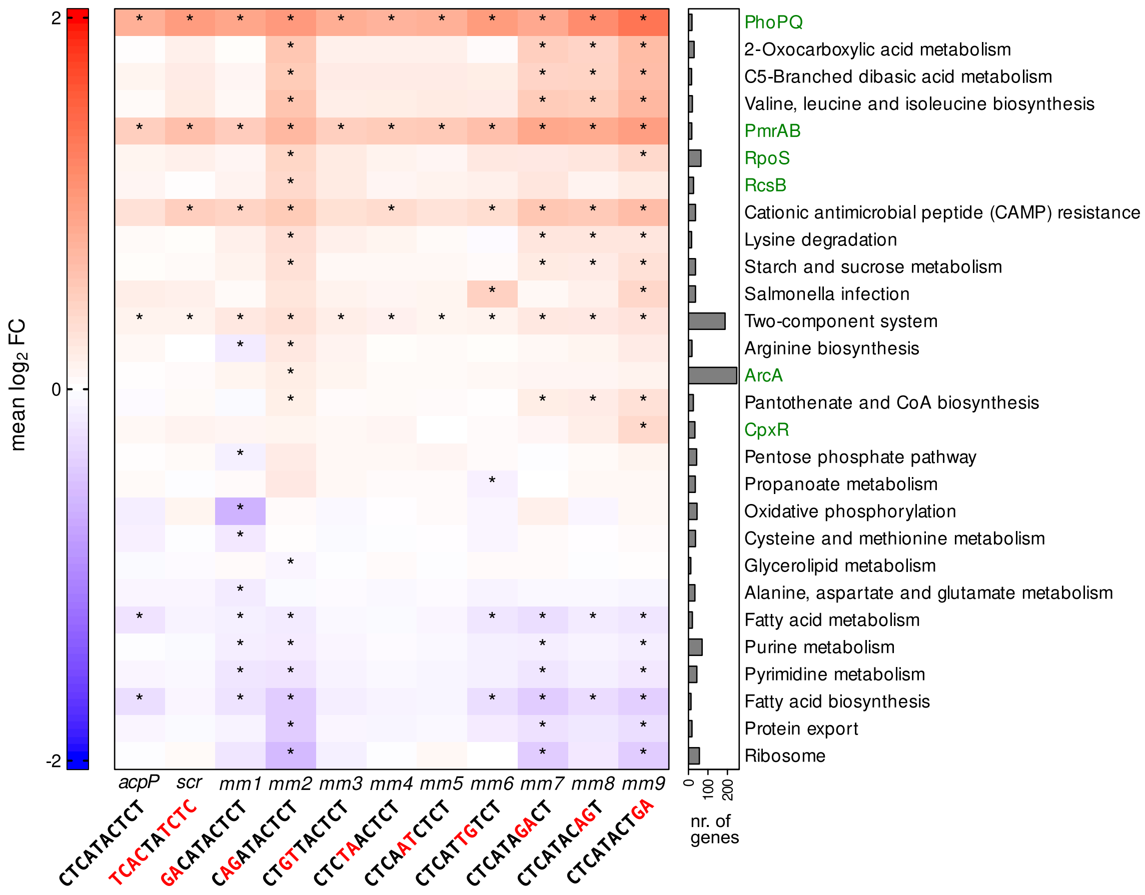


**Figure S9.** Gene set analysis reveals overall response to PNAs with differing sequences. Heatmap shows gene enrichment analysis classified by annotated KEGG pathways and known regulons (marked in green). Heatmap color shows the mean $\log_2$ fold change of genes belonging to the respective pathway or regulon. All samples were compared to the untreated control. Asterisks indicate a FRY gene set test FDR corrected p-value below 0.01 for the respective gene set. Shown are the 10 most significantly affected gene sets per condition. Barplot next to the heatmaps shows number of genes assigned to each gene set.

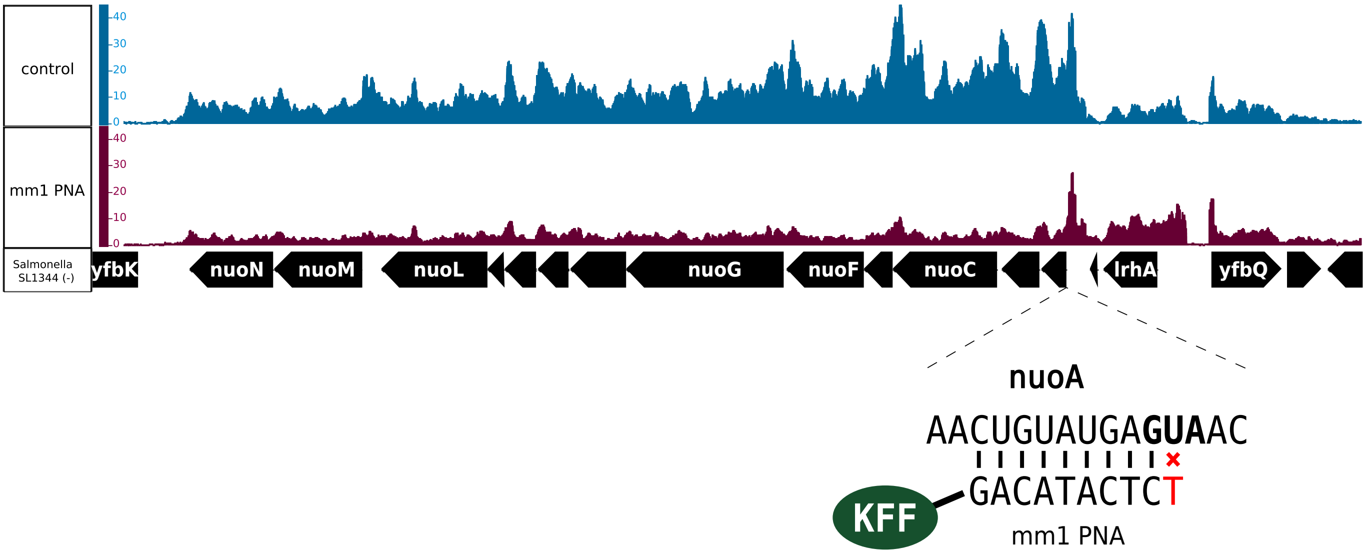


**Figure S10.** PNA affects multiple genes in downstream genes of the nuo operon. Read depth normalized by bins per million (BPM) are shown for PNA mm1 and the control is shown for comparison. *nuoA* is targeted by the mm1 PNA with one mismatch in its terminal position. The plots were created using the Affymetrix Integrated Genome Browser (IGB).

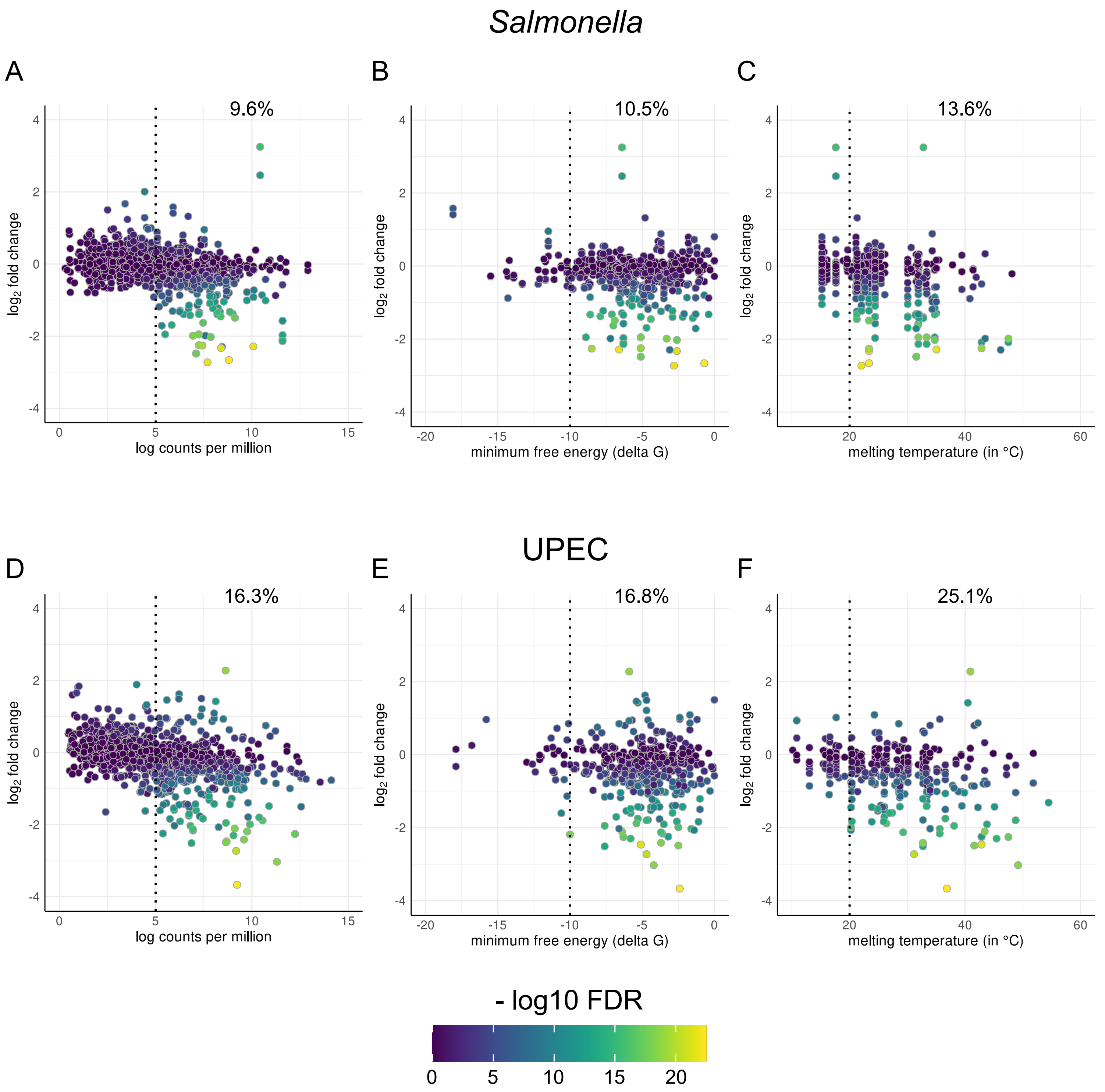


**Figure S11.** Transcript abundance, mRNA structure and PNA/mRNA melting temperature are related to off-target transcript's fold change. $Log_2$ fold changes of off-target transcripts of all PNAs tested with 0-3 mismatches and >6 consecutive matches were plottet against several factors. First, transcript abundance of the *Salmonella* (A) and UPEC (D) datasets were plotted and a threshold of 5 log CPM was set to separate transcripts with low average abundances. Next, minimum free energy of the remaning off-target mRNAs in the translation initiation region (-30 to +15 nt from start codon; B, E) were plotted and a threshold of -10 ΔG was set to separate TIRs with a strong predicted secondary structure. Finally, the predicted PNA/mRNA melting temperature was plotted (C, F) and a threshold of 20 °C was set to separate off-targets with low PNA/RNA binding affinity. FDR-corrected p-values are visualized by the fill color for the points. Percentages on the upper right of each plot denote the fraction of differentially depleted off-target genes after removal of transcripts below the respective threshold.

**Table S1.** PNA sequences used throughout this study. All PNAs were conjugated to the KFF peptide. Designed PNAs for both experiments with numbers of off-target sites inside translation initiation regions (TIRs) and total off-targets to all genes of *Salmonella enterica* serovar Typhimurium strain SL1344, as predicted by MASON. *N to C terminus orientation, # 5′ to 3′ orientation.

| PNA custom ID | PNA name | PNA sequence* | mRNA target sequence# | 0 mm off-targets | 0-3 mm off-targets (total) | 0-3 mm off-targets (TIR) | Minimal inhibitory concentration (MIC, in µM) | $OD_{600}$ (24h with 10 µM) |
|---|---|---|---|---|---|---|---|---|
| JVpna-14 | scrambled | TCACTATCTC | - | - | 1789 | 44 | >10 | 0.92 |
| *JVpna-15* | *acpP* | CTCATACTCT | | - | 494 | 16 | 1.25 | 0.00 |
| *Jvpna-29* | *acpP* mm1 | GACATACTCT | | - | 525 | 21 | 5 | 0.01 |
| *Jvpna-30* | *acpP* mm2 | CAGATACTCT | | - | 495 | 18 | >10 | 0.32 |
| *Jvpna-31* | *acpP* mm3 | CTGTTACTCT | | - | 1511 | 24 | >10 | 1.07 |
| *Jvpna-32* | *acpP* mm4 | CTCTAACTCT | AGAGU**AUG**AG | - | 811 | 23 | >10 | 0.89 |
| *Jvpna-33* | *acpP* mm5 | CTCAATCTCT | | - | 768 | 17 | >10 | 0.73 |
| *Jvpna-34* | *acpP* mm6 | CTCATTGTCT | | *ydhD* | 1566 | 31 | >10 | 0.46 |
| *Jvpna-35* | *acpP* mm7 | CTCATAGACT | | *kup* | 533 | 15 | >10 | 0.51 |
| *Jvpna-36* | *acpP* mm8 | CTCATACAGT | | - | 495 | 30 | 10 | 0.04 |
| *Jvpna-37* | *acpP* mm9 | CTCATACTGA | | - | 685 | 33 | 5 | 0.01 |
